# Mesenchymal stem cells ameliorate inflammation in an experimental model of Crohn’s disease via the mesentery

**DOI:** 10.1101/2023.05.22.541829

**Authors:** Maneesh Dave, Atul Dev, Rodrigo A Somoza, Nan Zhao, Satish Viswanath, Pooja Rani Mina, Prathyush Chirra, Verena Carola Obmann, Ganapati H Mahabeleshwar, Paola Menghini, Blythe Durbin Johnson, Jan Nolta, Christopher Soto, Abdullah Osme, Lam T Khuat, William Murphy, Arnold I Caplan, Fabio Cominelli

## Abstract

**Objective:** Mesenchymal stem cells (MSCs) are novel therapeutics for treatment of Crohn’s disease. However, their mechanism of action is unclear, especially in disease-relevant chronic models of inflammation. Thus, we used SAMP-1/YitFc, a chronic and spontaneous murine model of small intestinal inflammation, to study the therapeutic effect and mechanism of human bone marrow-derived MSCs (hMSC).

**Design:** hMSC immunosuppressive potential was evaluated through in vitro mixed lymphocyte reaction, ELISA, macrophage co-culture, and RT-qPCR. Therapeutic efficacy and mechanism in SAMP were studied by stereomicroscopy, histopathology, MRI radiomics, flow cytometry, RT-qPCR, small animal imaging, and single-cell RNA sequencing (Sc-RNAseq).

**Results:** hMSC dose-dependently inhibited naïve T lymphocyte proliferation in MLR via PGE_2_ secretion and reprogrammed macrophages to an anti-inflammatory phenotype. hMSC promoted mucosal healing and immunologic response early after administration in SAMP model of chronic small intestinal inflammation when live hMSCs are present (until day 9) and resulted in complete response characterized by mucosal, histological, immunologic, and radiological healing by day 28 when no live hMSCs are present. hMSC mediate their effect via modulation of T cells and macrophages in the mesentery and mesenteric lymph nodes (mLN). Sc-RNAseq confirmed the anti-inflammatory phenotype of macrophages and identified macrophage efferocytosis of apoptotic hMSCs as a mechanism of action that explains their long-term efficacy.

**Conclusion:** hMSCs result in healing and tissue regeneration in a chronic model of small intestinal inflammation. Despite being short-lived, exert long-term effects via macrophage reprogramming to an anti-inflammatory phenotype.

**Data Transparency Statement:** Single-cell RNA transcriptome datasets are deposited in an online open access repository ‘Figshare’ (DOI: https://doi.org/10.6084/m9.figshare.21453936.v1)

## INTRODUCTION

Crohn’s disease (CD) affects more than 1 million individuals in the US and is becoming more common worldwide.^1–3^ CD is thought to result from an inappropriate response of the host’s immune system to intestinal microbes.^4^ Therefore, the existing management strategies target inflammation and include immunosuppressive therapy with corticosteroids, immunomodulators, monoclonal antibodies against cytokines, and anti-adhesion molecules.^5^ However, only a fraction of patients started on these medications achieve and remain in remission for a long duration. In addition, the use of these immunosuppressive medications is associated with a risk of adverse events such as severe infections, and malignancies.^6^ Thus, there is a need for novel therapies that induce sustained remission with minimal side effects to treat CD. Cell-based therapies that utilize the immunosuppressive capacity of adult mesenchymal stem cells (MSCs) for immune-mediated diseases like CD are in clinical trials.^7–11^ A phase III multicenter randomized placebo-controlled trial of allogeneic adipose stem cells showed clinical and statistically significant healing of Crohn’s perianal fistulas and lead to its approval in European Union, Israel and Japan.^12, 13^ MSCs have good safety profile in clinical trials which has been attributed to their rapid clearance from the body after administration and yet have long term efficacy which is currently unexplained.^14, 15^ Furthermore, mechanistic studies in relevant and representative preclinical murine models of CD, especially chronic models are lacking. All the studies performed demonstrate the benefit of MSC therapy in murine models have been performed in only acute colitis models. The SAMP1/YitFc (SAMP) mice represent a unique model that spontaneously develops chronic small intestinal (SI) inflammation in the absence of genetic, chemical and immunological manipulation.^16^ This model also responds to conventional human CD therapies like anti-tumor necrosis factor-alpha agents^17^ and dexamethasone.^18^ Thus, the SAMP mice, which closely mimics human CD, can be utilized as a high-fidelity preclinical model for CD therapies, therefore we investigated efficacy and mechanism of human bone marrow derived MSC (hMSC) therapy in SAMP mice.

## METHODS

Standard methods (Single-cell RNA sequencing (Sc-RNA seq), RT-qPCR, enzyme immunoassay, mixed lymphocyte reaction (MLR), T cell proliferation assay using 3H-thymidine incorporation, flow cytometry, isolation of fibroblasts and hMSC, hMSC in vitro expansion and differentiation and additional details are described in the Supplementary Methods.

### Animal Experiments

Animal experiments were performed following the National Institutes of Health Guide for the Care and Use of Laboratory Animals. Protocols were approved by the Institutional Animal Care and Use Committee of the University of California, Davis (IACUC protocol # 21298) and Case Western Reserve University (CWRU; IACUC#2015-0142).

### hMSC treatment in SAMP

Isolation and expansion of MSC are described in supplementary methods. SAMP-1/YitFc mice with established inflammation (>25 weeks) were used for the in vitro and in vivo studies. In vitro cultured hMSC (5.0x10^6^) were resuspended in 200µl PBS and intraperitoneally injected in mice. An equivalent dose of 200 µl PBS served as vehicle control, while a daily dose of dexamethasone (DEX) (2mg/kg of mice weight) for seven days was used as a positive control.

### In vivo tracking of hMSC in SAMP and MRI

In vivo distribution of hMSC was performed using bioluminescent and NIR dye IVISense680 tagged hMSC. MRI was performed at the time of euthanasia in a 9.4 Tesla preclinical MRI scanner using the T2 fat suppression protocol. Complete experimental details of in vivo imaging and MRI are provided in supplementary methods.

### Single cell RNA sequencing

Single cell RNA sequencing was performed using SPLiTseq method from Parse Biosciences on single cell suspension from stromal vascular fraction (SVF) of mice mesentery. 20,000 cells were sequenced using NovaSeq PE100 illumine platform at the sequencing depth of 50000 reads/cells. Log fold change >1.5 and FDR<0.05 considered statistically significant in differentially expressed genes (DEGs).

### Data Analysis and Statistics

All the animals were randomized. No statistical method was used to predetermine the sample size. The sample size of each experiment and statistical details are provided in the respective figure legend. The statistical analysis was performed using GraphPad Prism (version 9.0; GraphPad Software, San Diego, CA). The student’s two-tailed unpaired t-test or Mann-Whitney test was used to compare the mean of two normally distributed groups. A one-way ANOVA test with Tukey as a posthoc test was applied for comparison between more than two groups. Data were expressed as mean ± SD. P values were considered statistically significant if *P<0.05, **P<0.01, ***P<0.001, each data point corresponds to a single mouse.

## RESULTS

### hMSCs suppress T cell proliferation in mixed lymphocyte reaction (MLR) and reprogram macrophages to an immunosuppressive phenotype

Human MSCs identified by expression of cell surface markers (CD73, CD90, CD105) and lack of hematopoietic marker CD34, CD14, HLA-DR **(Figure S1A)** have progenitor cell capacities that can be activated during in-vitro cell culture to differentiate into multi lineages **(Figure 1A);** human fibroblast served as study control and failed to differentiate **(Figure 1B)**. In addition to multilineage differentiation potential MSCs have immunosuppressive properties.^19^ We investigated the immunosuppressive potential of hMSC in mixed lymphocyte reaction (MLR) and macrophage co-culture assay using immune cells from SAMP. In MLR, co-culturing of mixed cultures of lymphocytes (SAMP splenocytes stimulated by irradiated allogeneic, C57Bl/6J derived splenocytes) with irradiated hMSC resulted in a dose-dependent inhibition of SAMP T-lymphocyte proliferation as measured by [3H] thymidine incorporation (P<0.0001); human dermal fibroblasts served as cell therapy control and did not suppress T cell proliferation (P=0.2699) **(Figure 1C).** Our previous study using interstitial cells of cajal stem cells identified prostaglandin E2 (PGE2) as an important mediator of immunosuppression,^20^ therefore, we measured PGE_2_ secretion by hMSC and in MLR supernatants using enzyme immunoassay. hMSC basally secrete PGE_2_ and the supernatants from suppressed T cells co-cultured with hMSC contained higher amounts of PGE_2_ than proliferating T cells cultured without hMSC **(Figure 1D).** Indomethacin, (COX-1 and 2 inhibitor) treated hMSC rescued T cell proliferation in MLR compared to hMSC (P<0.0001 **(Figure 1E).** Furthermore, we confirmed a significant reduction in PGE_2_ concentration in indomethacin treated hMSC groups **(Figure 1F).** Using the cells from MLR, we studied the expression of genes involved in Crohn’s disease pathogenesis using Crohn’s RT-PCR profiler array. hMSC treatment resulted in downregulation of 73 out of 84 pro-inflammatory CD genes (log2 fold change < 2; Q<0.05) **(Figure 1G).**

**Figure 1.**
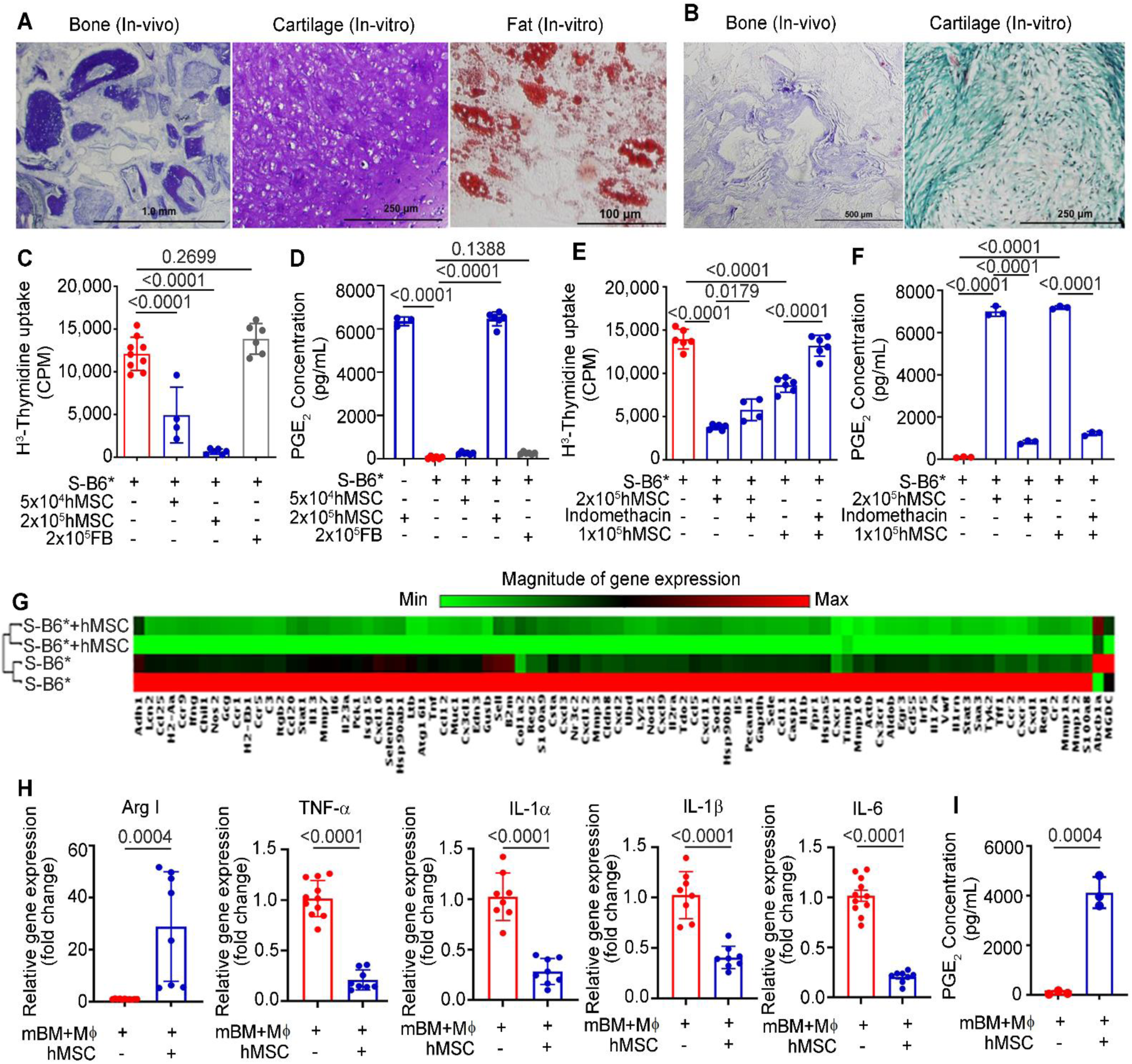
hMSC suppresses murine T cell proliferation and reprogram murine macrophages to an anti-inflammatory phenotype. **(A)** Mesodermal differentiation potential of BM-hMSCs and **(B)** Human fibroblasts (FB). **(C)** SAMP T cell proliferation in MLR (S: SAMP splenocytes, B6: C57Bl/6J derived irradiated splenocytes) treated with hMSC and FB for 96 hours. **(D)** PGE_2_ concentration measured in mixed lymphocyte reaction (MLR) supernatants using an enzyme immunoassay. **(E)** SAMP T cell proliferation in MLR treated with hMSC and COX inhibitor indomethacin (10µM) **(F)** PGE_2_ concentration measured in MLR supernatant using an enzyme immunoassay. **(G)** Heat Map showing the gene expression of Crohn’s RT-PCR profiler array on MLR reaction, **(H)** Relative gene expression of indicated markers and cytokine measured in total RNA extracted from murine bone marrow-derived macrophages (mBM-Mϕ). The gene expression was determined by qRT-PCR, normalized to GAPDH, and expressed as fold change (2-^ΔΔCt^). **(I)** PGE_2_ concentration measured in reaction supernatant from hMSC and murine bone marrow-derived macrophages (mBM-Mϕ) co-culture using an enzyme immunoassay. Data were expressed as mean ± SD from at least 2 independent experiments. *P<0.05, **P<0.01, ***P<0.001, by 2 tailed, unpaired Student’s t test.

PGE2 has been shown to promote polarization of macrophages to an anti-inflammatory phenotype, therefore we next performed hMSC co-cultures with bone marrow derived macrophages from SAMP (mBM- Mϕ). In cell proximity co-culture, where hMSC are in direct contact with macrophages, hMSC educated mBM-Mϕ to an anti-inflammatory phenotype with increased gene expression of Arginase I (28.9-fold change, P=0.0004), an anti-inflammatory phenotype marker and decreased gene expression of proinflammatory cytokines TNF-α (0.2-fold change, P<0.0001), IL-6 (0.2-fold change, P<0.0001), IL-1α (0.3-fold change, P<0.0001), and IL-1β (0.4-fold change, P<0.0001) **(Figure 1H).** Like MLR, the supernatants from mBM-Mϕ + hMSC group had significantly higher concentration of PGE2 (4100 ± 394 pg/mL; P=0.0004) compared to mBM-Mϕ **(Figure 1I)**. However, in transwell co-culture assay the separation of hMSC resulted in modest effect on macrophages with increased gene expression of Arginase-I (3.98-fold change, P<0.0001) and decrease in gene expression of TNF-α (0.64-fold change, P<0.001), and no significant changes in IL-6, IL-12 gene expression **(Figure S1B)**.

### Intraperitoneal hMSCs did not home to small intestine in SAMP

Previous studies in model of colitis have shown that intraperitoneal administration of MSCs can ameliorate colitis, therefore we next studied the homing potential and survival of hMSC after intraperitoneal administration in SAMP mice. The hMSCs were transduced with lentivirus to express firefly luciferase and in vivo animal bioluminescent imaging (BLI) was performed to detect live hMSCs in diseased SAMP and non-disease control AKR mice. We observed a time course decline in the BLI signal intensity of bioluminescent hMSCs with a drop in signal intensity on day 5 followed by a weak detectable signal on day 9 **(Figure 2A);** no signal was observed post day 9. Ex vivo imaging performed on harvested body organs on day 9 showed a weak BLI signal (live hMSC) in proximity of stomach in SAMP with no evidence of hMSC in the small intestine **(Figure S2).** To further confirm BLI data, we labeled hMSC with near-infrared fluorescent surface dye and performed time course epifluorescence imaging in diseased SAMP **(Figure 2B).** A similar trend of signal intensity was observed in epifluorescence imaging in SAMP, with a significant drop in signal intensity on day 9 (3.6e9±0.29e9, P=0.0403). Interestingly, we observed a very weak signal/hMSC presence in the peritoneum cavity on day 28 (0.49e9±0.44e9, P=0.0018) **(Figure 2C)** which is likely from remnant of dead hMSC as no live cells were observed by BLI or flow cytometry. We also performed ex vivo imaging of the peritoneum cavity in hMSC administered SAMP and found a strong signal of IVISense680 tagged hMSC on day 9 **(Figure 2D)** and a weak signal on day 28. We next performed flow cytometry on peritoneal lavage cells and demonstrated presence of live hMSCs (CD105^+^, CD73^+^) on day 9 **(Figure 2E)**, however, no viable hMSCs were detected on day 28. Furthermore, flow cytometry performed on single cell suspension from small intestine on day 9 and day 28 did not show presence of hMSC **(Figure S3**), confirming that majority of hMSCs do not migrate to small intestine.

**Figure 2.**
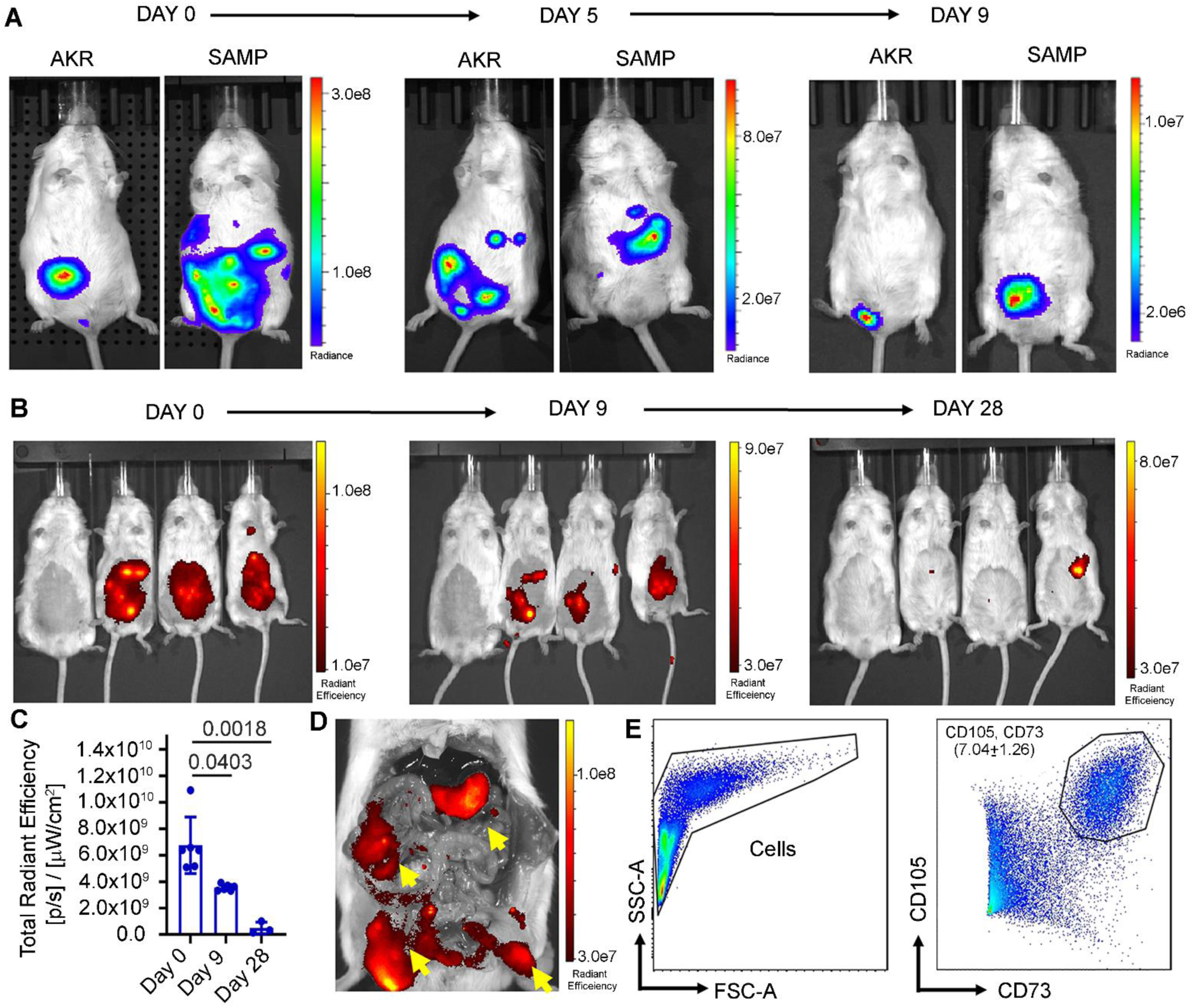
hMSC survive in the SAMP peritoneum cavity up to day 9. **(A)** Representative images showing time course BLI of transduced hMSC in AKR and SAMP mice on days 0, 5 and 9 **(B)** Representative time course epifluorescence images of SAMP mice showing the presence of intraperitoneally administered NIR dye IVIS680 labeled hMSCs. Images were acquired on days 0, 9 and day 28. **(C)** Quantitative estimation of total radiant efficiency in mice treated with IVIS680 tagged hMSCs. Data were expressed as mean ± SD from at least 2 independent experiments. *P<0.05, **P<0.01, ***P<0.001, by 1-way ANOVA. **(D)** Representative ex-vivo epifluorescence image of SAMP peritoneum cavity showing the presence of IVIS680 tagged hMSCs (yellow arrow) on day 9. **(E)** Flow cytometry gating scheme to detect hMSC cell population in the peritoneal lavage of hMSC administered SAMP at day 9.

### hMSC result in mucosal healing and immunologic response early after administration

As intraperitoneally injected live hMSCs population were detected only till day 9, we studied their therapeutic response in SAMP mice with established chronic inflammation at day 9 after hMSC treatment. SAMP mice were treated with 5 million hMSC; high dose dexamethasone ^21^ served as the positive therapy control, while phosphate buffer saline (PBS) treated mice served as negative control. Mice in DEX group were injected with a daily dose of DEX for 7 days prior to the onset of the treatment regimen **(Figure 3A)**. 3D stereomicroscopy which quantifies spatial abnormalities **(Figure S4)** of the intestine^22^ showed mucosal healing with significant decrease in percent abnormal mucosa in hMSC treatment group than PBS (P=0.0065)(**Figure 3B, 3C).** Blinded histopathological scoring using a standardized, previously validated scoring system that employed for SAMP mice^23^ showed a partial response in the DEX group (P=0.0563) while no response was observed in hMSC group (0.8282)**(Figure S5).** hMSC and DEX treated SAMP showed some improvement in epithelial regeneration, however complete healing was not achieved. As our in vitro data showed hMSC induced T cell suppression, we studied the effect of hMSC treatment on T cell population in mesenteric lymph node (mLN). We observed a significant decrease in absolute numbers of CD3^+^ (P=0.0312) and CD4^+^ (0.0272) cells; no significant decrease (P=0.1155) observed in CD8^+^ cells **(Figure 3F, 3G).** Next, we studied gene expression of cytokines in mLN cells; hMSC treated mice showed decrease in gene expression of cytokines involved in Th-1 pathway (TNF-α; 0.60-fold change, P<0.0001); Th-2 pathway (IL-4; 0.54-fold change, P=0.0274), and Th-17 pathway and (IL-21; 0.28-fold change, P<0.0001). **(Figure 3H).** We further investigated the extended effect of DEX treatment on established intestinal inflammation in SAMP till day 28 **(Figure 3I)**. SAMP mice treated with DEX show non-significant (P=0.9631) changes in percentage of abnormal mucosa compared to PBS treated group **(Figure 3J, K).** Blinded histopathological scoring showed severe disease recurrence at day 28 with non-significant changes (P=0.4079) in inflammatory index between PBS and DEX treated SAMP **(Figure 3L, M).**

**Figure 3.**
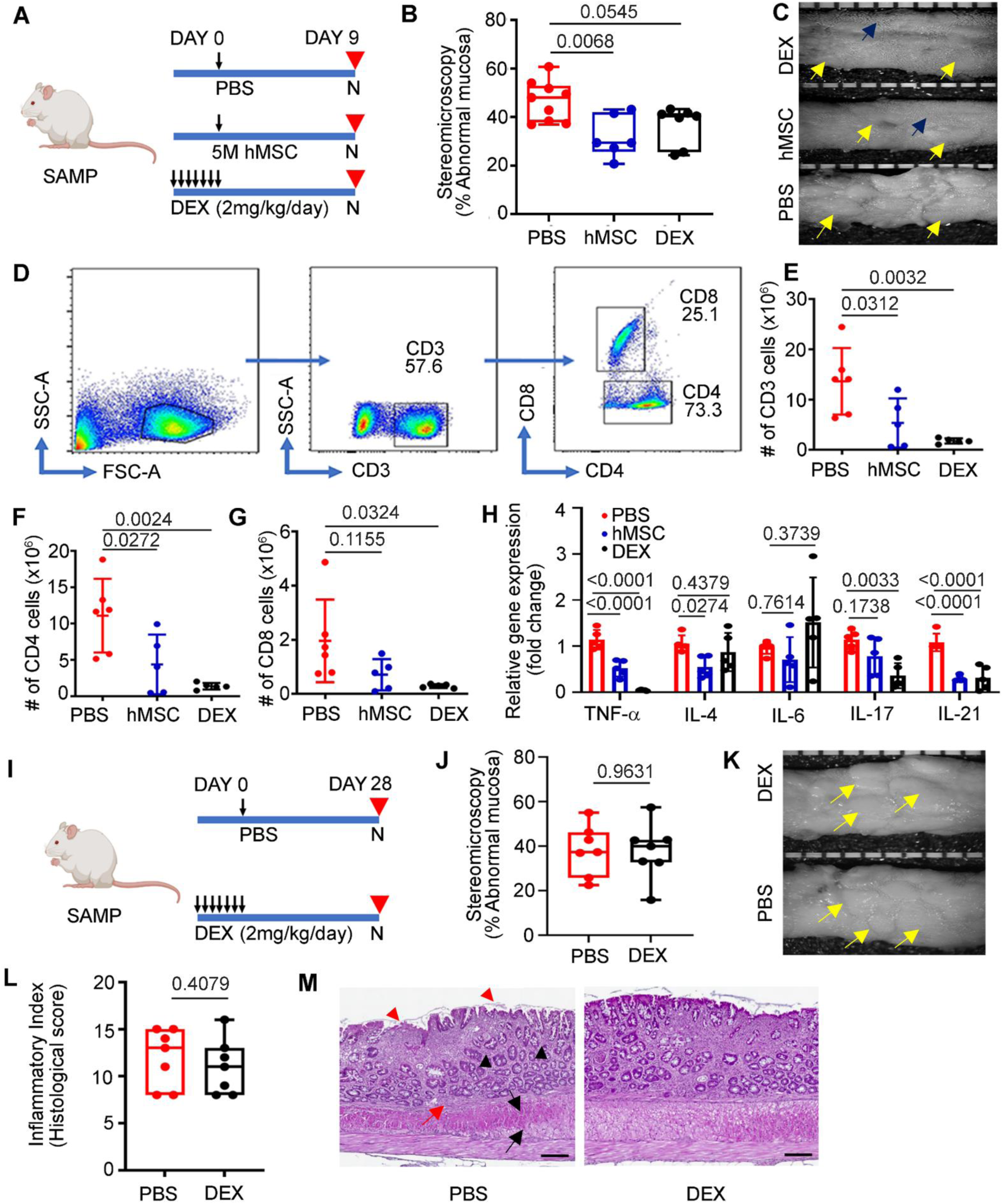
hMSC treatment shows mucosal healing and early sign of immunological response on day 9. **(A)** Schematic showing the treatment regimen of the experiment (M=million, N=Necropsy). **(B)** Percent abnormal mucosa in PBS, hMSCs and DEX-treated groups. **(C)** Representative comparative stereoscopic view of small intestine from DEX, hMSCs and PBS-treated groups (yellow arrow showing abnormal mucosa; blue arrow showing normal mucosa). **(D)** Flow cytometry gating strategy showing **(E)** Absolute number of CD3^+^, **(F)** CD4^+^ and **(G)** CD8^+^ cells in single cell suspension of mesenteric lymph node (mLN). **(H)** Relative gene expression of cytokines from Th-1, Th-2 and Th-17 pathways was measured in total RNA extracted from mesenteric lymph node cells. Gene expression was determined by qRT-PCR, normalized to GAPDH, and expressed as fold change (2-ΔΔCt). **(I)** Schematic showing treatment regimen of the experiment **(J)** Percent abnormal mucosa in PBS and DEX treated groups. **(K)** Representative comparative stereoscopic view of the small intestine in PBS and DEX treated groups (yellow arrows showing abnormal mucosa **(L)** The inflammatory index for disease severity in SAMP mice treated with PBS and DEX **(M)** Representative histopathology photomicrograph of ileum tissue from PBS, and DEX treated groups (Red triangle= villus distortion, black triangle=crypts dysplasia, red arrow=immune infiltration in submucosa, black arrow= muscle hypertrophy). The scale bar represents 200µm. Data were expressed as mean ± SD from at least 2 independent experiments, each data point represent one mouse. *P<0.05, **P<0.01, ***P<0.001, by 1-way ANOVA and unpaired t-test **(J and L)**.

### hMSC alleviate established small intestinal inflammation in SAMP mice and result in mucosal, histological, and radiological healing by day 28

As there was an early but incomplete response after hMSC therapy, we next studied therapeutic response on day 28 of the treatment when no live hMSCs are present. High dose DEX administered daily for 7 days ameliorates inflammation, however the inflammation eventually recurs after stopping DEX. SAMP mice treated with DEX show severe disease recurrence at day 28 after treatment, therefore, for the day 28 experiment, we modified our DEX treatment approach (positive therapy control) and administered it daily for 7 days just prior to euthanasia **(Figure 4A).** In these experiments, SAMP mice were treated with 5 million hMSC; high dose dexamethasone served as the positive therapy control, while phosphate buffer saline (PBS) treated mice served as negative control. hMSC (PBS: 44.80± 15.57 vs. hMSC: 29.36± 20.29; P=0.0284) and DEX (PBS: 44.80± 15.57 vs. DEX: 14.17±4.56; P<0.0001) had mucosal healing as demonstrated by significant reduction in percent abnormal mucosa **(Figure 4B, 4C).** Inflammatory Index for disease severity is a histological index composed of the villus distortion index, active inflammation index, mononuclear inflammation index, chronic inflammation index and transmural inflammation index. SAMP mice treated with hMSC had reduced severity of small intestine (SI) inflammation (PBS: 18±3.55 vs. hMSC: 12.43±7.5; P=0.0171); SAMP mice treated with DEX (2 mg/kg/24hr i.p.) for 7 days prior to euthanasia had more reduction of SI inflammation as compared to the hMSC treatment (PBS: 18±3.55 vs. DEX: 5.18±2.18; P<0.001) **(Figure 4D, 4E)** manifesting in more intact villi and reduced immune infiltrate and transmural inflammation. Next, we studied the gene expression of cytokines in mLN cells; hMSC treated mice showed decrease in gene expression of pro-inflammatory cytokines from Th-1 pathway (TNF-α and IFN-γ); Th-2 pathway (IL-6, IL-4) and Th-17 pathway (IL-17, IL-21). DEX served as positive control for the treatment, and significantly downregulated gene expression of majority of proinflammatory cytokines including TNF-α, IFN-γ, IL-6, IL-17, IL-21 **(Figure S6).**

**Figure 4.**
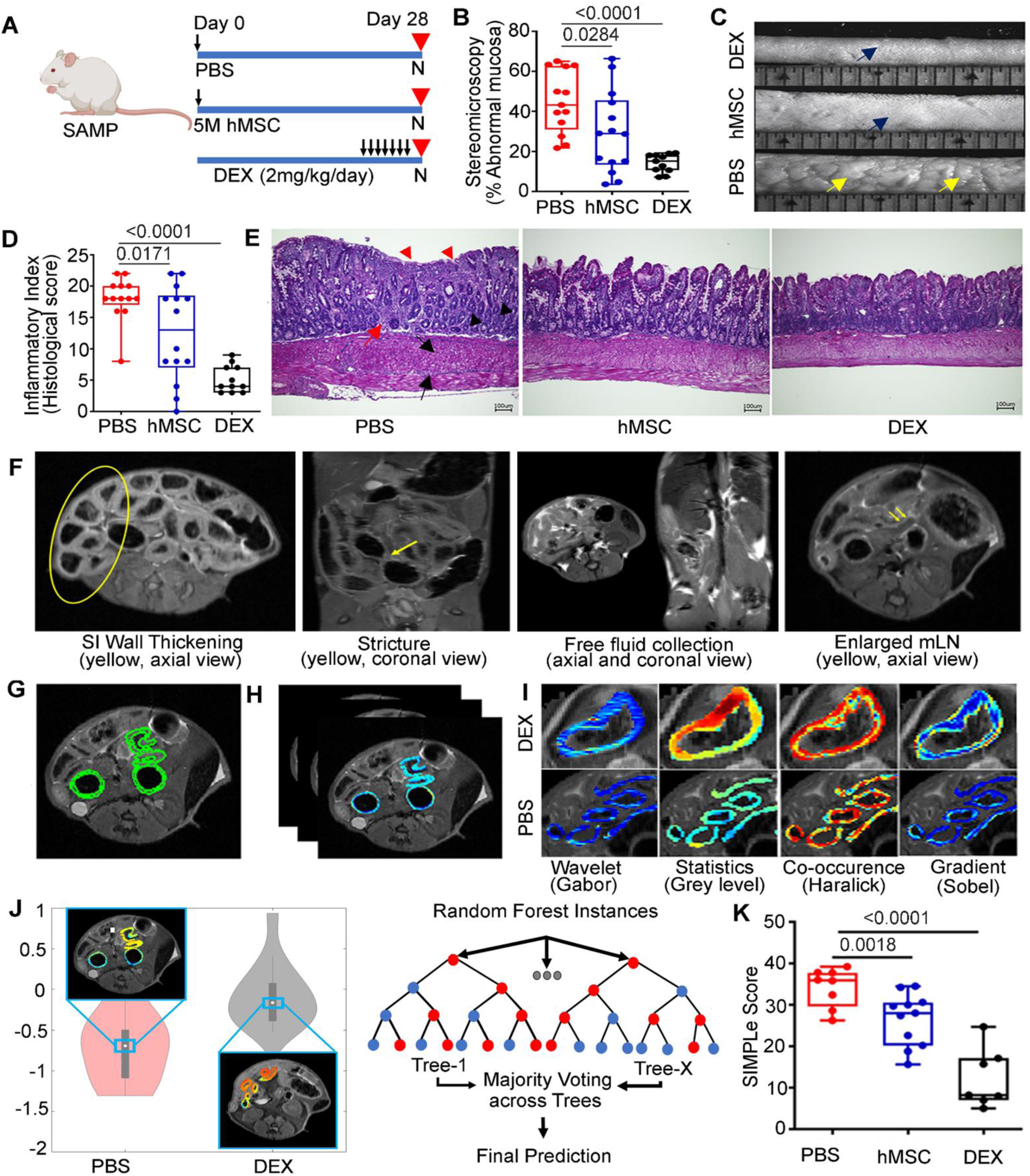
hMSC treatment alleviate intestinal inflammation in SAMP on day 28. **(A)** Schematic showing the treatment regimen of the experiment. **(B)** Percent abnormal mucosa in PBS, hMSCs and DEX-treated groups. **(C)** Representative comparative stereoscopic view of the small intestine in DEX, hMSCs and PBS-treated groups (yellow arrows showing abnormal mucosa; blue arrow showing normal mucosa). **(D)** The Inflammatory Index for disease severity in PBS, hMSCs and DEX-treated groups showing histologic small intestine (SI) inflammation **(E)** Representative histopathology photomicrograph of ileum tissue from PBS, hMSC and DEX-treated groups. The scale bar represents 200µm. **(F)** MR image for SAMP shows human CD features, including SI wall thickening, strictures, free fluid collection, enlarged mesenteric lymph node(mLN). **(G)** MR image annotation in SAMP, **(H)** Radiomic feature extraction **(I)** Radiomics analysis involved 100 radiomic features from 4 different classes to quantify SI wall appearance in terms of heterogeneity and gradient responses in PBS and DEX-treated groups. **(J**) Top-ranked features (Haralick and Sobel) were selected and used to train a machine learning Random Forest classifier to yield a radiomics-based likelihood of disease severity. **(K)** SIMPle score resulted in further enhancement of disease severity assessment and discrimination between treatment groups. Data were expressed as mean ± SD from at least 3 independent experiments, each data point represent one mouse. *P<0.05, **P<0.01, ***P<0.001, by 1-way ANOVA.

Radiologic healing in small bowel Crohn’s disease is associated with a reduction in the risk of poor outcomes in CD patients.^24^ We previously extracted radiomic features (quantitative computer-extracted image descriptions) from magnetic resonance imaging (MRI) of CD patients and showed their ability in combination with clinical phenotype to predict the risk of surgery.^25^ Therefore, we studied radiologic healing in SAMP mice using a new scoring method that incorporates objective MRI radiomic analysis and pathology to accurately characterize disease in the small intestine. SAMP mice with established SI inflammation were found to exhibit MRI features of human CD including SI wall thickening, strictures, free fluid collections, and enlarged mesenteric lymph nodes **(Figure 4F).** MRI images from SAMP were annotated by a blinded radiologist **(Figure 4G)** and radiomic features were extracted **(Figure 4H)**. Radiomic feature selection was performed using imaging samples from the DEX (low disease severity) and PBS (high disease severity) groups **(Figure 4I)**. Features with significant separation between these groups were identified utilizing Wilcoxon rank sum. The topmost discriminatory radiomic features were used to train a Random Forest predictive model which provided a probability of a high severity sample **(Figure 4J).** This model was then applied to the hMSCs treated and additional PBS and DEX treated groups. There was a positive correlation between MRI inflammatory scores and pathology scores **(Figure S6).** Combining radiomics score with pathology score (SIMPle score) resulted in further enhancement of disease severity assessment and discrimination between groups, with hMSC treated mice demonstrating radiologic healing (PBS:34.2±1.6 vs hMSC:26.5±1.9; ANOVA P=0.0018, PBS:34.2±1.6 vs DEX: 12.2±2.7; ANOVA P<0.0001) **(Figure 4K).** Overall, our results demonstrate that while dexamethasone is potent, its effect is transient, whereas hMSC have sustained effect at day 28.

### Single cell RNA sequencing of mesentery identified a novel macrophage pattern of hMSC mediated response

Our imaging data showed hMSC on the mesentery of SAMP mice **(Figure 2D)** and given increasing recognition of the importance of mesenteric inflammation in the pathophysiology of Crohn’s disease, we performed Sc-RNAseq on single cell suspension from the stromal vascular fraction (SVF) of mice mesentery on day 9 and day 28 of hMSC treatment using SPLiT-seq method. Raw sequencing data was processed to qualify the quality criteria **(Figure S7).** Transcripts from mice treated with hMSC at day 9 clustered into 10 groups at clustering resolution 1.0 with variable number of cells and UMI counts per cell **(Figure 5A).** These clusters were annotated to fibroblast, macrophages, T Lymphocytes, epithelial cells, mesothelial cells, smooth muscle cells and Schwan cells using scMRMA and after manual curing based on available literature on cell types in Sc-RNAseq. Differential expressed gene (DEG) analysis performed on all cell types revealed highly enriched population of differentially expressed genes in hMSC treated mice (FDR<0.05) with five genes (*Camk1d, Lars2, Xist, Gphn, Gm19951*) reaching statistical significance **(Figure 5B).** As macrophage constitutes a major component of the mesentery and our in vitro data on macrophage polarization to anti-inflammatory phenotype, we studied in-depth the transcriptomic profile of the macrophage cluster. We observed larger numbers of differentially expressed genes in macrophage cluster in hMSC treated mice with 33 genes meeting statistical significance, among which, *Lars2, Cmss1, Gphn*, were top three downregulated genes while *Map4, Usp34,* and *Lrrfip 1* genes were upregulated after hMSC treatment **(Figure 5C).** *Mrc1,*^26^ *Cd163,*^27^ *H2-Ab1*^28^ are marker genes that are highly expressed in macrophages and were used to represent macrophage cluster in UMAP plot **(Figure 5D).** Based on published studies^29, 30^ and heterogeneity in macrophage phenotype, we further analyzed a set of genes (*Mrc1, Cd163, Basp1, Mgl2, Lvye1, Retnla, Adgre1, Apoe, F13a1*)^31–35^ that represents M1 pro-inflammatory and M2 anti-inflammatory phenotype of macrophages in different tissues. Analogous to our in vitro data, we observed a marked increase in the average gene expression of genes representing M2 anti-inflammatory macrophages *(Mrc1, Cd163, Mgl2, H2-Ab1* and *Retnla)* in hMSC treatment group, however the percentage of cells expressing these genes did not change across treatment **(Figure 5E).**

**Figure 5.**
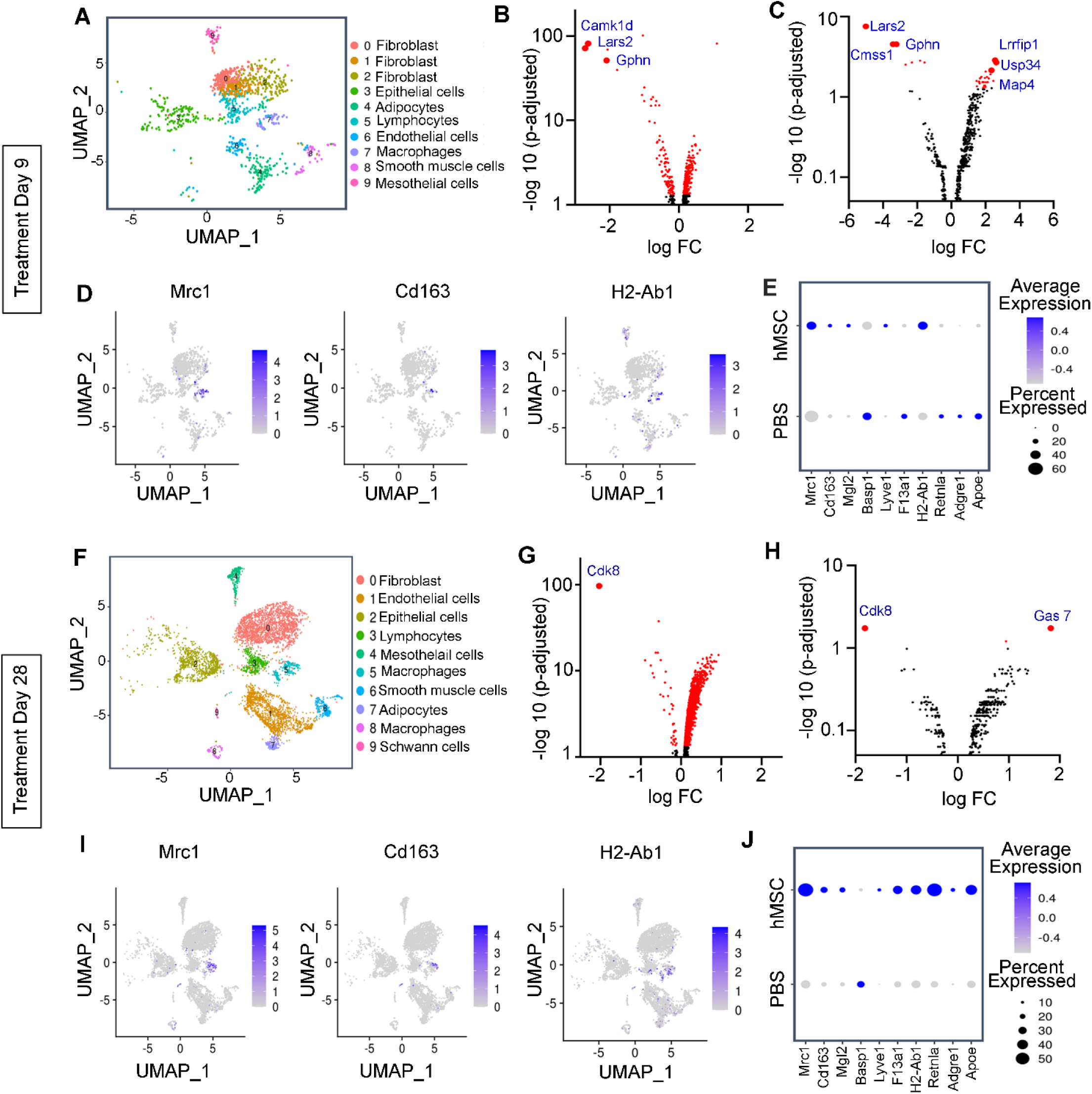
Sc-RNAseq was performed on single-cell suspension of mesenteric stromal vascular fraction (SVF) from SAMP mice treated with PBS and hMSC. **(A-E)** Sc-RNAseq showing analysis on day 9 samples. **(A)** UMAP plot showing identified clusters at a resolution of 1.0. **(B)** Volcano plots showing differential gene expression in all cell types with a threshold value (FDR<0.05). The top 3 significant differentially expressed genes (logFC >1.5 with FDR <0.05) are highlighted and labelled in the plots. **(C)** Volcano plots showing differential gene expression in macrophage cluster with a threshold value (FDR<0.05). The top 3 significant differentially expressed genes (logFC>1.5 with FDR <0.05) (upregulated and downregulated) are labelled and highlighted in the plot. **(D)** UMAP plots of the representative macrophage markers genes (*Mrc1, Cd163, H2-Ab1*) showing normalized gene expression in the macrophage cluster. **(E)** Dot plots showing the average expression of selective genes expressed by the proportion of the cells in the macrophage cluster across the treatment. **(F-J)** Single-cell RNA sequencing showing analysis on day 28 samples. **(F)** UMAP plot showing identified clusters at cluster resolution 0.4. **(G)** Volcano plots show differential gene expression in all cell types with a threshold value (FDR<0.05). Cdk8 was significant differentially expressed gene (logFC >1.5 with FDR <0.05) in hMSC-treated mice on day 28. **(H)** Volcano plots showing differential gene expression in macrophage cluster with a threshold value (FDR<0.05). Two genes were significantly differentially expressed (FC >1.5 with FDR <0.05) in hMSC-treated mice. **(I)** UMAP plots of the representative macrophage markers genes (*Mrc1, Cd163, H2-Ab1*) showing normalized gene expression in the macrophage cluster **(J)** Dot plots showing average expression of selective genes and proportion of the cells representing in the cluster across the treatment.

Similarly, we have analyzed the therapeutic effect of hMSC treatment at day 28 on the cellular composition and transcriptomic alterations in all detected cell types. The detected transcripts from PBS and hMSC treated SAMP were clustered into 10 groups on UMAP plots **(Figure 5F)** and annotated to fibroblast, macrophages, T Lymphocytes, epithelial cells, mesothelial cells, smooth muscle cells and schwann cells using scMRMA method and manually cured. DEG analysis on all cell types revealed highly enriched population of differentially expressed genes in hMSC group, with *Cdk8* gene demonstrating statistical significance **(Figure 5G).** DEG analysis in macrophage cluster revealed several DEGs in hMSC group, with downregulated Cdk8 (kinase involved in inflammatory responses) and upregulated Gas7 (gene involved in phagocytic cup formation in macrophage) genes met statistical significance criteria **(Figure 5H).** Similar to the UMAP plot of day 9 samples, *Mrc1, Cd163, H2-Ab1* marker genes represent macrophage population on UMAP plots on day 28 samples **(Figure 5I).** Lastly, we studied average gene expression and percent representation of cells expressing set of genes in macrophage cluster using dot plot analysis. We found that hMSC treatment resulted in pronounced increase in the average gene expression of anti-inflammatory M2 phenotype *(Mrc1, Cd163, Mgl2, Lyve1, F13a1, Retnla, and Apoe)* along with an increase in anti-inflammatory macrophage numbers compared to the early response on day 9 **(Figure 5J).**

### hMSC educate macrophages to anti-inflammatory phenotype and suppress T cell proliferation to alleviate small intestine inflammation in SAMP

We further confirmed immunomodulatory effect of hMSC on CD11b^+^ macrophages isolated from mLN, mesentery and peritoneal macrophages at day 9 and day 28 after hMSC treatment. At day 9, hMSC treatment resulted in anti-inflammatory phenotype of mLN and SVF macrophages with decreased gene expression of proinflammatory cytokines TNF-α and increase in M2 anti-inflammatory phenotype marker Arginase-I **(Figure 6A, 6B).** CD11b^+^ macrophages isolated from mLN and SVF of mesentery at day 28 also demonstrated similar anti-inflammatory phenotype with increase in gene expression of surface marker Arginase-I and decrease in gene expression of proinflammatory cytokines TNF-α **(Figure 6C, 6D).** Additionally, we studied gene expression of several other proinflammatory cytokines including IL-1α, IL-1β, IL-12 and IL-6 at day 9 and day 28 of the treatment and most were downregulated after hMSC treatment **(Figure S8).** Furthermore, we performed flow-based cell sorting of peritoneal macrophages Gr1, F4/80 and demonstrated that peritoneal macrophages from SAMP mice treated with hMSCs had an anti-inflammatory phenotype with higher Arginase-I/TNF-α ratio (P=0.0261) **(Figure 6E).** As no live hMSC were detected at day 28 and given upregulation of Gas7, a marker of phagocytosis at day 28, we next assessed the ability of SAMP macrophages to phagocytose hMSCs via efferocytosis and therefore acquire anti-inflammatory phenotype.^36^ Hence, we injected celltracker Red^TM^ CMTPX dye labelled hMSC in SAMP and noted that CD11b+ macrophages efferocytose apoptotic hMSCs in peritoneal cavity and mesenteric SVF **(Figure 6F, 6G)** and these macrophages have anti-inflammatory phenotype with significantly higher Arginase-I/TNF-α ratio (P=0.0037) **(Figure 6H)**. Furthermore, our in vitro co-culture assay with live and apoptotic hMSCs demonstrated that SAMP macrophages did not show active efferocytosis with live hMSC while significantly uptake apoptotic hMSCs with visible formation of phagosome. We confirmed our results by labeling apoptotic hMSCs with annexin to mark compromised cell membrane, and these marked apoptotic hMSC were detected in the F4/80 tagged peritoneal macrophages on co-culture **(Figure 6I)**.

**Figure 6.**
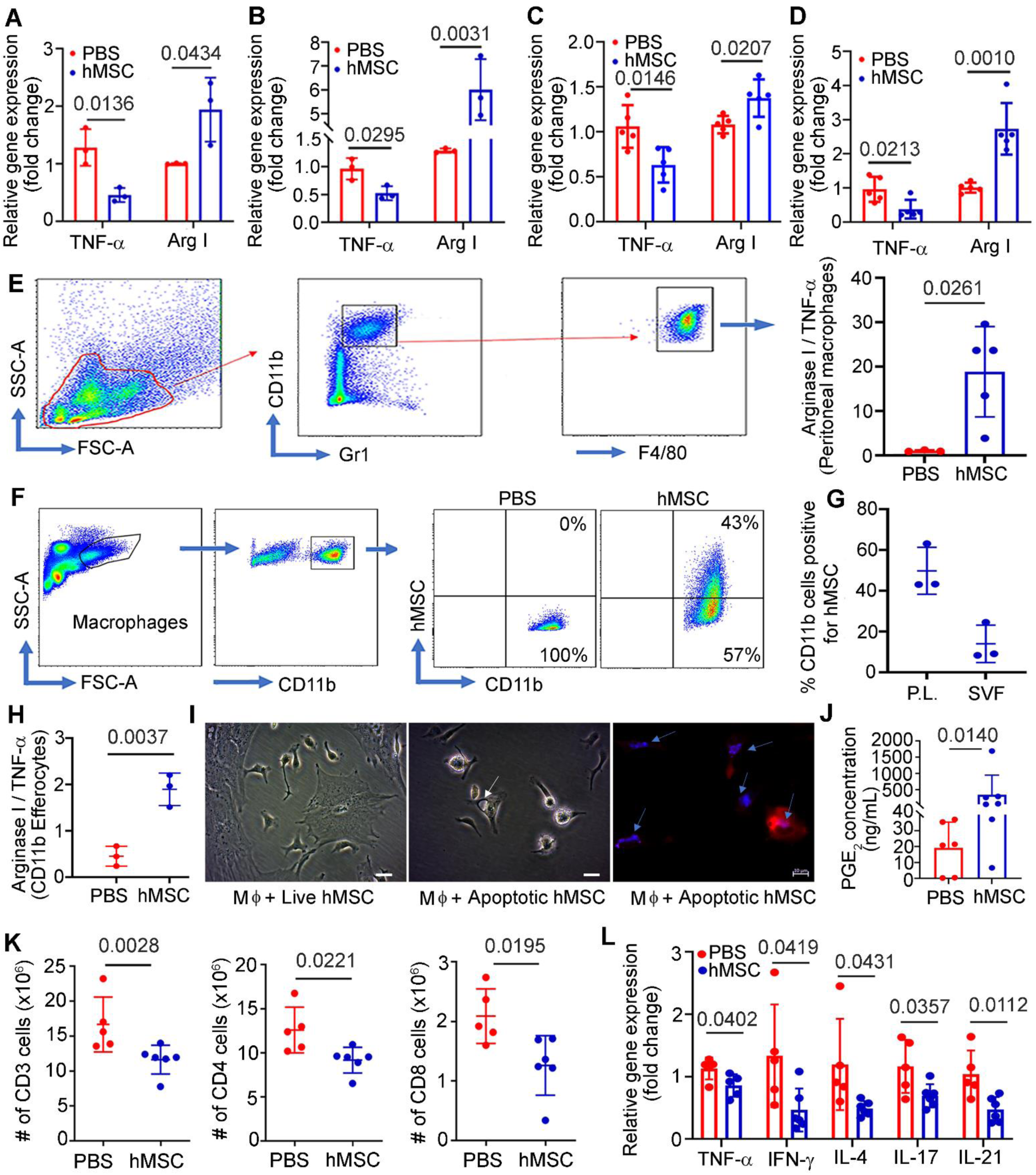
hMSC secrete PGE_2_, educates mLN, mesenteric and peritoneum macrophages to anti-inflammatory phenotype, and suppress T cells to alleviate small intestine inflammation in SAMP. **(A, B)** Relative gene expression of TNF-α and Arginase I on day 9 of the treatment, measured in the total RNA extracted from CD11b^+^ cells from **(A)** mesenteric lymph node and **(B)** mesenteric stromal vascular fraction. **(C, D**) Relative gene expression of TNF-α and Arginase I on day 28 of the treatment measured in the total RNA extracted from CD11b^+^ cells from **(C)** mesenteric lymph node **(D)** mesenteric stromal vascular fraction. Gene expression was determined by qRT-PCR, normalized to GAPDH, and expressed as fold change (2-ΔΔCt). **(E)** Arginase 1/ TNF-α ratio measured by flow cytometry in SAMP peritoneal macrophages. **(F, G)** Cell Tracker Red^TM^ CMPTX dye labelled hMSC showing in-vivo phagocytosis by CD11b^+^ macrophages in peritoneal lavage (P.L.) and SVF at day 9. **(H)** Arginase 1/ TNF-α ratio measured by flow cytometry in SAMP peritoneal Cd11b^+^ macrophages (Efferocytes) phagocytosed hMSC **(I)** Brightfield microscopic images of SAMP peritoneal macrophages co-cultured with live hMSCs (no visible apoptosis observed) and apoptotic hMSCs (white arrow showing phagosome formation in macrophage), fluorescence image showing phagocytosis of apoptotic hMSC by SAMP peritoneal macrophages, arrows showing engulfed apoptotic bodies of hMSC. Scale bar 10 µm. **(J)** PGE_2_ concentration measured in the peritoneal fluid of SAMP. **(K)** Absolute numbers of CD3, CD4, and CD8 lymphocytes measured in mLN at day 28 of the treatment. **(L)** Relative gene expression of cytokines in CD4 lymphocytes from hMSC-treated SAMP at day 28. Data were expressed as mean ± SD from at least 2 independent experiments, each data point represent one mouse. *P<0.05, **P<0.01, ***P<0.001, by 2 tailed, unpaired Student’s t test.

In addition to efferocytosis, we detected significantly higher concentration of PGE_2_ (341 ± 650 ng/mL; P=0.0140) in SAMP peritoneal fluid after intraperitoneal injection of hMSC **(Figure 6J).** Similar to our in-vitro data which demonstrated suppression of T cell proliferation by PGE_2_ secretion by hMSCs, we found that hMSC resulted in T cell suppression in mLN with significant decrease in absolute numbers of CD3(PBS:16.65±3.91; hMSC:11.63±2.06, P=0.0228), CD4(PBS:12.59±2.60, hMSC:9.165±1.45, P=0.0221) and CD8(PBS: 2.088±0.45, hMSC:1.261±0.50, P=0.0195) cells **(Figure 6K)** and immunosuppressive phenotype of CD4 cells with decrease in gene expression of pro-inflammatory cytokines from Th-1 (TNF-α ; 0.85-fold change, P=0.0402, IFN-γ; 0.46-fold change, P=0.0419), Th-2(IL-4; 0.49-fold change, P=0.0431), and Th-17 (IL-17;0.69-fold change, P=0.0357, IL-21;0.47-fold change, P=0.0112) pathways **(Figure 6L).** Briefly, hMSCs secrete PGE_2_ in peritoneal cavity and lymphatic vessels to suppress T cell proliferation and reprogram macrophages to anti-inflammatory phenotype and enhances macrophages efferocytosis capacity ^37^. Peritoneal and SVF macrophage perform efferocytosis of apoptotic hMSCs becoming polarized to anti-inflammatory phenotype followed by clonal expansion which results in long term efficacy **(Figure 7).**

**Figure 7.**
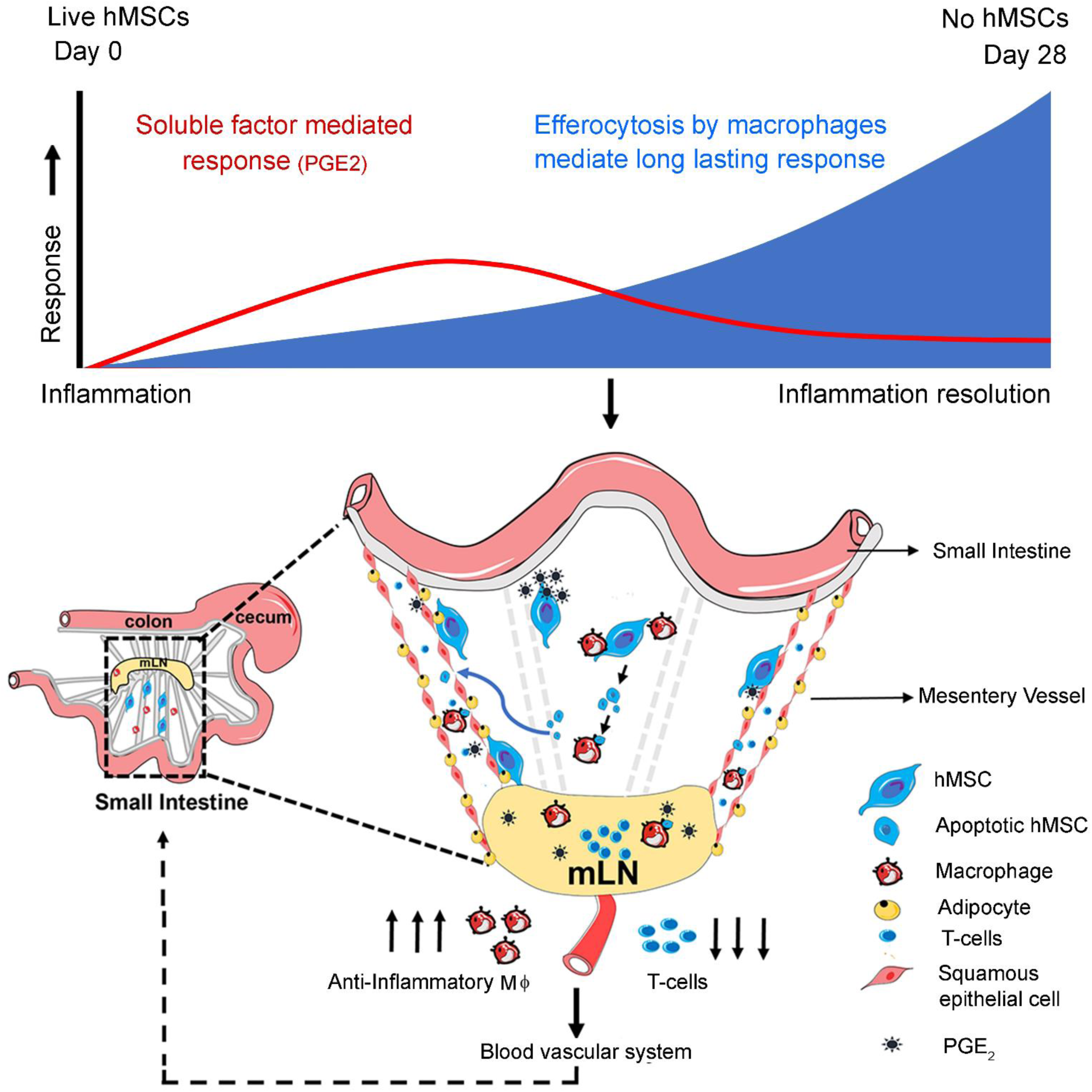
Schematic showing putative mechanism of hMSCs mediated immunosuppression in experimental model of human Crohn’s disease.

## DISCUSSION

MSCs are a novel therapeutic approved in EU, Japan, Israel for treatment of CD perianal fistulas and in advanced stage clinical trials for immune mediated diseases like CD,^7^ however, their mechanism of action in CD is unclear. Here we show that hMSCs have a potent immunosuppressive activity in vitro and result in histologic, mucosal and radiologic healing of chronic intestinal inflammation in a spontaneous model of human Crohn’s disease. Following intraperitoneal administration, hMSC did not home to small intestine and mediated their effect via modulation of T cells and macrophages in the mesentery and mesenteric lymph node. Furthermore, our results demonstrate that while a steroid like dexamethasone is potent, its effect is transient, whereas hMSC have prolonged effect that outlast their presence.

Using multiple imaging modalities and flow cytometry we demonstrated that after intraperitoneal injection majority of hMSCs did not migrate to the small intestine and remained in the peritoneal cavity adjacent to the mesentery and lymph node. Our data is analogous to previous studies that show limited homing of MSCs to the colon after intraperitoneal administration in colitis models.^38^ As a very small fraction of hMSC survived beyond day 9 of administration, we studied hMSC therapeutic effect and mechanism at day 9 and noted mucosal healing and immunological response. Since most hMSCs did not migrate to small intestine, their secretory product including PGE_2_^39^ likely diffused through the fenestrated lymphatic vessels^40, 41^ to access mesentery and reach mLN through the lymphatic drainage. These secretory products suppressed T cell proliferation, educated macrophages to anti-inflammatory phenotype, and this was supported by several lines of in-vitro and in-vivo evidence including; hMSC secretion of PGE_2_, reduction of hMSC effect on T-cell response by pharmacological blockade of PGE_2_, higher concentration of PGE_2_ in SAMP peritoneal fluid after intraperitoneal injection of hMSC, reduction in absolute numbers of CD3^+^ and CD4^+^ T cells in SAMP mLN with downregulation of gene expression of Th-1 and Th-2 cytokine, and conversion of macrophages to an anti-inflammatory phenotype in-vitro and in mLN, mesenteric and peritoneum macrophages of SAMP, and macrophage efferocytosis. As mesentery plays an important role in pathogenesis of Crohn’s disease,^42–44^ we further characterized hMSC therapeutic mechanism by unbiased ScRNA seq on SVF mesenteric cells and noted hMSC treatment resulted in differential expression of genes across all detected cell types in SVF. Mesenteric inflammation is implicated in Crohn’s disease however, no medical therapy specifically targets mesenteric inflammation. Our study shows that mesentery could be targeted for the treatment of small bowel inflammation.

ScRNA seq provided further evidence of macrophage polarization to anti-inflammatory phenotype as hMSC downregulated the expression of several genes involved in inflammatory pathways in macrophages. hMSC treatment upregulated gene expression of transcriptional repressor *Lrrfip1* in macrophages, which codes for the Leucine-rich repeat flightless-interacting protein 1 and directly bind to the GC-rich sequences of the tumor necrosis factor-alpha (TNF-α) promoter and inhibit its production.^45^ We also observed an increase in the average expression of a set of genes (*Mrc1, Cd163, Mgl2, Lyve1, F13a1, Retnla, and Apoe*), representing heterogeneous tissue resident anti-inflammatory M2 macrophages in the mesentery of hMSC treated SAMP, however number of these macrophages did not change across treatment groups by day 9. At day 28, hMSC treatment results in a complete response in SAMP mice marked by mucosal, histological, immunologic, and radiological healing. Immunologically, there was reduction in absolute numbers of CD3^+^, CD4^+^ T and CD8^+^ T cells in SAMP mLN with downregulation of gene expression of multiple Th-1 and Th-2 cytokines, and increase in anti-inflammatory phenotype in mLN, mesenteric and peritoneum macrophages of SAMP treated with hMSC. The increased time required for a complete response is likely due to the significant histologic inflammation in SAMP mice with established disease that require longer time to heal as well as the unique mechanism of action of hMSCs. Our day 28 ScRNA seq data showed downregulation of Cdk8 and upregulation of Gas7 in macrophages and demonstrated an increase in the number of anti-inflammatory macrophages compared to day 9. Interestingly, these genes were not DEGs in macrophages during the early phase, indicating the dominance of this mechanism in the mesenteric macrophages in the absence of live hMSC, which could explain their anti-inflammatory phenotype. Gas7 is involved in development of phagocytic cap and is highly expressed in macrophages capable of phagocytosis.^46^ Efferocytosis, a process of clearing dead and apoptotic bodies by phagocytic cells, can induce proliferation in macrophages and result in an anti-inflammatory phenotype.^36^ Our data shows that SAMP macrophages phagocytose hMSCs in peritoneal cavity and result in anti-inflammatory phenotype; a significant percentage of hMSC apoptotic bodies are uptaken by SVF resident macrophages which resulted in a marked increase in numbers of heterogeneous SVF resident M2 macrophages with higher expression for *Mrc1, Cd163, Mgl2, Lyve1, F13a1, Retnla,* and *Apoe* genes at day 28. Our data is congruent to a study that used clinical samples from patients with graft-versus-host-disease treated with MSCs and murine models to show that MSCs undergo apoptosis after infusion, and this is the primary driver of their immunosuppressive function and clinical response in patients.^47^

In summary, MSCs are short lived after administration and yet can have prolonged therapeutic effects which have been unexplained. Our results demonstrate that intraperitoneally administered hMSC modulates the immune response in SAMP in stages. In the early stage, live hMSC secretion of molecules like PGE_2_ is the dominant mechanism. These molecules released by hMSCs in the peritoneal cavity diffuse through the fenestrated mesenteric lymphatic vessels to reach mLN, inhibit naïve T cell proliferation and educate SVF and mLN resident macrophages to anti-inflammatory phenotype with subsequent downregulation of proinflammatory cytokines, and enhance macrophages efferocytosis capacity. Secretory molecules like PGE_2_ work universally and induce immunosuppression across the species as demonstrated by our in vitro MLR and macrophage co-culture data. In the later phases following hMSC treatment, macrophages efferocytose apoptotic hMSCs which lead to their proliferation and a reprogramming to anti-inflammatory phenotype which is maintained and mediates the long-term immune effects. Meriwether et al showed that after efferocytosis, macrophages upregulate Cox2 and PGE_2_ production which had been previously shown to act via EP2 receptors on macrophages.^37, 48^ This then led to increased activity of Rac1, which resulted in increases in the numbers of macrophages efferocytosing apoptotic cells. ^48^ Metabolites produced from apoptotic cells after efferocytosis have been demonstrated to reprogram macrophages in multiple ways^49, 50^. Yurdagul Jr et al demonstrated that arginine produced in macrophages after efferocytosis promotes continual efferocytosis via HuR protein mediated stabilization of *Mcf2* mRNA that facilitates Rac1 activation and leads to resolution of injury in an atherosclerosis model ^50^. Furthermore, Ampomah et al demonstrated that apoptotic cell-derived methionine induces the activity of DNA methyltransferase 3A (DNMT3A) that induces epigenetic modification that represses *DUSP4* function, allowing for uninterrupted ERK signaling and production of PGE_2_ and TGFβ.^49^ In our study, these anti-inflammatory macrophages suppress the proliferation and numbers of potentially pathogenic T cells in mLN resulting in reduced numbers in the small intestine contributing to the marked reduction in small intestine inflammation.

The study has a few limitations, including the employment of human MSCs in an immune dysregulated mouse. Human and mouse MSCs are considered immune-evasive and characteristically have low expression of MHC II, CD40, CD80 and CD86^51, 52^ and multiple prior studies have utilized human MSCs to ameliorate inflammation in murine models of acute colitis and did not find any induction of immunological response.^15, 53^ In addition, the mechanism of action that demonstrates PGE_2_ is an important molecule that induces immunosuppression across the species; and macrophage phagocytosis of apoptotic hMSC partly obviates these concerns. Another limitation is that a small percentage of MSCs undetectable by our current methods could have localized to the small intestine, however that likely has limited role in therapeutic efficacy.

hMSC are short lived which contributes to their safety profile noted in human clinical studies but exert immunosuppressive and tissue regenerative properties that outlast their presence in the body, our study provides a mechanism for this observation. Mesenteric inflammation is implicated in Crohn’s disease however, no medical therapy specifically targets mesenteric inflammation. Our study shows that mesentery could be targeted for the treatment of small bowel inflammation via MSCs and shows multiple potential avenues for enhancing the potency of MSCs via modulation of their secretome/PGE2 and/or their efferocytosis ability, which can then be tested for efficacy in SAMP.

## ACKNOWLEDGEMENTS

This work was supported by Crohn’s and Colitis Foundation Career Development Award 370615 (to M.D.) NIH grants: K08DK110421 (to M.D.), F.C. supported by P30DK097948, DK042191, DK091222 and, DK07948 grants, A.I.C. supported by P41 EB021911 grant, G.H.M. supported by Crohn’s and Colitis Foundation Senior Research Award 421904, HL126626 and HL141423 grants, V.C.O supported by Swiss national science foundation grant: P2SKP3_178132. The authors thank Margie Harris for the assistance with hMSC isolation and culture, Mike Sorrell for isolation of human dermal fibroblast. We duly acknowledge The DNA Technologies and Expression Analysis Core, Bioinformatics Core, Genome Center UC Davis, Dr. Lutz Froenicke, Dr. Hong Qiu, and Dr. Jie Li, for single cell RNA sequencing. We also thank Dr. Chris A. Flask, Case Center for Imaging Research, Case Western Reserve University and Center for Molecular and Genomic Imaging, UC Davis for providing small animal in vivo imaging facilities.

## DECLARATION OF INTERESTS

The Authors have no interests to declare.

## INCLUSION AND DIVERSITY

We support inclusive, diverse, and equitable conduct of research.

## Notes

https://doi.org/10.6084/m9.figshare.21453936.v1

### Competing Interest Statement

The authors have declared no competing interest.

## REFERENCES

1. Molodecky NA, et al. Increasing incidence and prevalence of the inflammatory bowel diseases with time, based on systematic review. Gastroenterology 142, 46–54 e42; quiz e30 (2012).

2. Burisch J, Jess T, Martinato M, Lakatos PL, EpiCom E. The burden of inflammatory bowel disease in Europe. Journal of Crohn’s & colitis 7, 322–337 (2013).

3. Collaborators GBDIBD. The global, regional, and national burden of inflammatory bowel disease in 195 countries and territories, 1990-2017: a systematic analysis for the Global Burden of Disease Study 2017. Lancet Gastroenterol Hepatol 5, 17–30 (2020).

4. Abraham C, Cho JH. Inflammatory bowel disease. The New England journal of medicine 361, 2066–2078 (2009).

5. Baumgart DC, Le Berre C. Newer Biologic and Small-Molecule Therapies for Inflammatory Bowel Disease. The New England journal of medicine 385, 1302–1315 (2021).

6. Cohn HM, Dave M, Loftus EV, Jr. Understanding the Cautions and Contraindications of Immunomodulator and Biologic Therapies for Use in Inflammatory Bowel Disease. Inflamm Bowel Dis 23, 1301–1315 (2017).

7. Ko JZ, Johnson S, Dave M. Efficacy and Safety of Mesenchymal Stem/Stromal Cell Therapy for Inflammatory Bowel Diseases: An Up-to-Date Systematic Review. Biomolecules 11, (2021).

8. Garcia-Olmo D, Garcia-Arranz M, Herreros D, Pascual I, Peiro C, Rodriguez-Montes JA. A phase I clinical trial of the treatment of Crohn’s fistula by adipose mesenchymal stem cell transplantation. Diseases of the colon and rectum 48, 1416–1423 (2005).

9. Duijvestein M, et al. Autologous bone marrow-derived mesenchymal stromal cell treatment for refractory luminal Crohn’s disease: results of a phase I study. Gut 59, 1662–1669 (2010).

10. Liang J, et al. Allogeneic mesenchymal stem cell transplantation in seven patients with refractory inflammatory bowel disease. Gut 61, 468–469 (2012).

11. Ciccocioppo R, et al. Autologous bone marrow-derived mesenchymal stromal cells in the treatment of fistulising Crohn’s disease. Gut 60, 788–798 (2011).

12. Panes J, et al. Expanded allogeneic adipose-derived mesenchymal stem cells (Cx601) for complex perianal fistulas in Crohn’s disease: a phase 3 randomised, double-blind controlled trial. Lancet 388, 1281–1290 (2016).

13. Panes J, et al. Long-term Efficacy and Safety of Stem Cell Therapy (Cx601) for Complex Perianal Fistulas in Patients With Crohn’s Disease. Gastroenterology 154, 1334–1342 e1334 (2018).

14. Lalu MM, et al. Safety of cell therapy with mesenchymal stromal cells (SafeCell): a systematic review and meta-analysis of clinical trials. PloS one 7, e47559 (2012).

15. Dave M, Jaiswal P, Cominelli F. Mesenchymal stem/stromal cell therapy for inflammatory bowel disease: an updated review with maintenance of remission. Curr Opin Gastroenterol 33, 59–68 (2017).

16. Pizarro TT, et al. SAMP1/YitFc mouse strain: a spontaneous model of Crohn’s disease-like ileitis. Inflamm Bowel Dis 17, 2566–2584 (2011).

17. Marini M, et al. TNF-alpha neutralization ameliorates the severity of murine Crohn’s-like ileitis by abrogation of intestinal epithelial cell apoptosis. Proc Natl Acad Sci U S A 100, 8366–8371 (2003).

18. McNamee EN, et al. Novel model of TH2-polarized chronic ileitis: the SAMP1 mouse. Inflamm Bowel Dis 16, 743–752 (2010).

19. Viswanathan S, et al. Mesenchymal stem versus stromal cells: International Society for Cell & Gene Therapy (ISCT(R)) Mesenchymal Stromal Cell committee position statement on nomenclature. Cytotherapy 21, 1019–1024 (2019).

20. Dave M, et al. Stem cells for murine interstitial cells of cajal suppress cellular immunity and colitis via prostaglandin E2 secretion. Gastroenterology 148, 978–990 (2015).

21. Stoll BJ, et al. Trends in Care Practices, Morbidity, and Mortality of Extremely Preterm Neonates, 1993-2012. Jama 314, 1039-1051 (2015).

22. Rodriguez-Palacios A, et al. Stereomicroscopic 3D-pattern profiling of murine and human intestinal inflammation reveals unique structural phenotypes. Nature communications 6, 7577 (2015).

23. Burns RC, Rivera-Nieves J, Moskaluk CA, Matsumoto S, Cominelli F, Ley K. Antibody blockade of ICAM-1 and VCAM-1 ameliorates inflammation in the SAMP-1/Yit adoptive transfer model of Crohn’s disease in mice. Gastroenterology 121, 1428–1436 (2001).

24. Deepak P, et al. Radiological Response Is Associated With Better Long-Term Outcomes and Is a Potential Treatment Target in Patients With Small Bowel Crohn’s Disease. The American journal of gastroenterology 111, 997–1006 (2016).

25. Chirra P, et al. Integrating Radiomics With Clinicoradiological Scoring Can Predict High-Risk Patients Who Need Surgery in Crohn’s Disease: A Pilot Study. Inflamm Bowel Dis, (2022).

26. Jablonski KA, et al. Novel Markers to Delineate Murine M1 and M2 Macrophages. PloS one 10, e0145342 (2015).

27. Hu JM, et al. CD163 as a marker of M2 macrophage, contribute to predicte aggressiveness and prognosis of Kazakh esophageal squamous cell carcinoma. Oncotarget 8, 21526–21538 (2017).

28. Stables MJ, et al. Transcriptomic analyses of murine resolution-phase macrophages. Blood 118, e192–208 (2011).

29. Tabula Sapiens C, et al. The Tabula Sapiens: A multiple-organ, single-cell transcriptomic atlas of humans. Science 376, eabl4896 (2022).

30. Dominguez Conde C, et al. Cross-tissue immune cell analysis reveals tissue-specific features in humans. Science 376, eabl5197 (2022).

31. van Kooyk Y, Ilarregui JM, van Vliet SJ. Novel insights into the immunomodulatory role of the dendritic cell and macrophage-expressed C-type lectin MGL. Immunobiology 220, 185–192 (2015).

32. Kieu TQ*, et al*. Kinetics of LYVE-1-positive M2-like macrophages in developing and repairing dental pulp in vivo and their pro-angiogenic activity in vitro. Scientific reports 12, 5176 (2022).

33. Lee MR*, et al*. The adipokine Retnla modulates cholesterol homeostasis in hyperlipidemic mice. Nature communications 5, 4410 (2014).

34. Waddell LA, et al. ADGRE1 (EMR1, F4/80) Is a Rapidly-Evolving Gene Expressed in Mammalian Monocyte-Macrophages. Frontiers in immunology 9, 2246 (2018).

35. Gafencu AV, Robciuc MR, Fuior E, Zannis VI, Kardassis D, Simionescu M. Inflammatory signaling pathways regulating ApoE gene expression in macrophages. The Journal of biological chemistry 282, 21776–21785 (2007).

36. Mehrotra P, Ravichandran KS. Drugging the efferocytosis process: concepts and opportunities. Nature reviews Drug discovery 21, 601–620 (2022).

37. Frasch SC, et al. Signaling via macrophage G2A enhances efferocytosis of dying neutrophils by augmentation of Rac activity. The Journal of biological chemistry 286, 12108–12122 (2011).

38. Sala E, et al. Mesenchymal Stem Cells Reduce Colitis in Mice via Release of TSG6, Independently of Their Localization to the Intestine. Gastroenterology 149, 163–176 e120 (2015).

39. Kalinski P. Regulation of immune responses by prostaglandin E2. Journal of immunology 188, 21–28 (2012).

40. Sabine A, Davis MJ, Bovay E, Petrova TV. Characterization of Mouse Mesenteric Lymphatic Valve Structure and Function. Methods in molecular biology 1846, 97–129 (2018).

41. Murfee WL, Rappleye JW, Ceballos M, Schmid-Schonbein GW. Discontinuous expression of endothelial cell adhesion molecules along initial lymphatic vessels in mesentery: the primary valve structure. Lymphat Res Biol 5, 81–89 (2007).

42. He Z, et al. Microbiota in mesenteric adipose tissue from Crohn’s disease promote colitis in mice. Microbiome 9, 228 (2021).

43. Coffey JC, O’Leary DP, Kiernan MG, Faul P. The mesentery in Crohn’s disease: friend or foe? Curr Opin Gastroenterol 32, 267–273 (2016).

44. Peyrin-Biroulet L, et al. Mesenteric fat as a source of C reactive protein and as a target for bacterial translocation in Crohn’s disease. Gut 61, 78–85 (2012).

45. Suriano AR, et al. GCF2/LRRFIP1 represses tumor necrosis factor alpha expression. Molecular and cellular biology 25, 9073–9081 (2005).

46. Hanawa-Suetsugu K, et al. Phagocytosis is mediated by two-dimensional assemblies of the F-BAR protein GAS7. Nature communications 10, 4763 (2019).

47. Galleu A, et al. Apoptosis in mesenchymal stromal cells induces in vivo recipient-mediated immunomodulation. Science translational medicine 9, (2017).

48. Meriwether D, et al. Macrophage COX2 Mediates Efferocytosis, Resolution Reprogramming, and Intestinal Epithelial Repair. Cellular and molecular gastroenterology and hepatology 13, 1095–1120 (2022).

49. Ampomah PB, et al. Macrophages use apoptotic cell-derived methionine and DNMT3A during efferocytosis to promote tissue resolution. Nat Metab 4, 444–457 (2022).

50. Yurdagul A, Jr.*, et al.* Macrophage Metabolism of Apoptotic Cell-Derived Arginine Promotes Continual Efferocytosis and Resolution of Injury. Cell metabolism 31, 518–533 e510 (2020).

51. Koppula PR, Chelluri LK, Polisetti N, Vemuganti GK. Histocompatibility testing of cultivated human bone marrow stromal cells - a promising step towards pre-clinical screening for allogeneic stem cell therapy. Cellular immunology 259, 61–65 (2009).

52. Krampera M, Galipeau J, Shi Y, Tarte K, Sensebe L, Therapy MSCCotISfC. Immunological characterization of multipotent mesenchymal stromal cells--The International Society for Cellular Therapy (ISCT) working proposal. Cytotherapy 15, 1054–1061 (2013).

53. Duijvestein M*, et al*. Pretreatment with interferon-gamma enhances the therapeutic activity of mesenchymal stromal cells in animal models of colitis. Stem cells 29, 1549–1558 (2011).

