## Supplementary information for "Mesenchymal stem cells ameliorate inflammation in an experimental model of Crohn’s disease via the mesentery"

**Conflict-of-interest**: The authors have declared that no conflict of interest exists.

### SUPPLEMENTARY METHODS

#### Mice

SAMP1/YitFc (SAMP) mice were obtained from Case Western Reserve University (CWRU) under the material transfer agreement and rederived at UC Davis. Initial experiments on SAMP mice were performed at Case Western Reserve University (CWRU; IACUC#2015-0142) and then with SAMP mice housed at UC Davis vivarium. The animals were housed in AAALAC approved specific pathogen-free facility of Institute of Regenerative cure, UC Davis Health, Sacramento and maintained in Tecniplast IVC (Individually Ventilated Cages) Blue line, following the standard 12-h light/dark cycles. All mice had ad libitum access to water and were fed with the standard global 18% protein rodent diet throughout the experiments. All experimental procedures were approved by the Institutional Animal Care and Use Committee of UC Davis (IACUC protocol # 21298). Both male and female mice with established disease (>24 weeks old) were randomly assigned to the control and treatment groups to minimize the confounders. All outcome assessments of the experiments including histopathology scoring, stereomicroscopy, RT qPCR, radiomics and MRI scoring were performed in blinded manner.

#### Bone marrow-derived human mesenchymal stem cells (BM-hMSCs) isolation and expansion

Bone marrow were collected from healthy de-identified donors; the procedure was reviewed and approved by the University Hospitals of Cleveland and University of California Davis Institutional Review Boards. The hMSCs were isolated as per previously described protocols[1, 2] and expanded in vitro in standard tissue culture vessels at $1.0 \times 10^4$ cells per $cm^2$. The hMSCs were successfully stored at P3 in liquid nitrogen and subsequently expanded in T175mm tissue culture flask using DMEM, selected lots of 10% FBS supplemented with 2mM L-glutamine and 1% penicillin-streptomycin before injection. The cells were maintained in the $CO_2$ incubator at 37°C and 5% $CO_2$ for 9-11 days, culture media was changed every 3-4 days.

**Fibroblasts isolation and expansion**:

Human skin fibroblasts were isolated as previously described method.[3] Briefly, Human skin, keratomed at a depth of 0.4 mm from the buttocks region of volunteers, was provided by the Skin Diseases Research Center, Case Western Reserve University School of Medicine, under an approved IRB protocol. The epidermal and dermal components were separated by treatment with 50 U/ml dispase (Collaborative Biomedical Products, Bedford, Mass.) overnight at 4°C. The dermal component was minced and incubated at 37°C for 1 h with an enzyme cocktail consisting of 100 mM sodium pyruvate, 0.1 mg/ml DNase, 1.25 mg/ml hyaluronidase, and 2.7 mg/ml collagenase C in DMEM-LG buffered with HEPES, pH 7.4. The resulting tissue suspension was filtered through 100-μm nylon meshes to separate single cells and small aggregates from larger pieces of tissue and cultured in DMEM-LG supplemented with 10% FBS.

**Chondrogenic differentiation:**

Cells were trypsinized and then resuspended in chondrogenic differentiation medium (DMEM-high glucose supplemented with 1% ITS+,$10^{-7}$ M dexamethasone, 1mM sodium pyruvate, 120mM ascorbic acid-2 phosphate, 100mM nonessential amino acids, and 10ng/mL TGF-β1). Two hundred microliters of this cell suspension containing 250,000 cells was added per well of a 96-well polypropylene V-bottom, multiwell dish (Phenix Research). The multiwell plates were centrifuged at 500 g for 5 min and then incubated at 37°C. The differentiation medium was changed every other day. Chondrogenic pellets were harvested after 21 days for histological analysis.

**In vivo osteogenesis:**

In vivo osteogenesis of MSCs is tested by their capability of forming ectopic bone tissue, Briefly, cells are vacuum-loaded into fibronectin-coated 3x3x3 mm hydroxyapatite/tricalcium phosphate cubes generously provided by the Zimmer Corporation (Warsaw, IN). Cell-loaded cubes were then incubated at 37C for 2 h in a humidified atmosphere of 5% CO2 and 95% air. After the incubation period, individual cubes were implanted subcutaneously on the dorsal surface of severe combined immunodeficient mice following an approved IACUC protocol. Animals were euthanized after 6 weeks, and the cubes were fixed with 10% phosphate-buffered formalin.

**Cell transduction:**

hMSC were seeded at $1\times10^4$ cells per $cm^2$ and transduced with a multiplicity of infection (MOI) of 5 in the presence of protamine sulfate and transduced for 24 hours. Three days after the last round of transduction, the efficiency is measured by detecting the RFP by flow cytometry. To evaluate that the transduction process does not affect the stem cell properties of MSCs, the chondrogenic (in vitro) and osteogenic (in vivo) capabilities were assessed.

**MLR reaction**

Splenocytes were isolated from two different strains of mice (C57BL/6J and SAMP/YitFc). One set of splenocytes served as responder cells while the other set was irradiated at 3300 Gy and served as stimulator cells. Briefly, $1x10^6$ activated mixed splenocytes [SAMP/YitFc(responders)+C57BL/6(stimulator)] were co-cultured with the irradiated human mesenchymal stem cells (3300 Rad for 10 min) in different cell ratios (1:20, 1:10,1:5) in microwell plates for 4 days. Human dermal Fibroblast was taken as a cell therapy control. The suppression of the T-cells was studied after adding the $^3$H-Thymidine on day 4. The plates were Incubated for another 18 hours in radioactive incubator during which $^3$H-thymidine will be incorporated into the newly synthesized DNA of the dividing cells. The proliferation was calculated measuring the uptake of [3H] thymidine in a scintillation counter and is expressed as counts per million.

### Macrophage and hMSC co-culture

SAMP bone marrow was harvested and enriched for CD14 cells using CD14 MACS purification kit. CD14 monocytes were further cultured in complete RPMI1640 with 50ng/ml MCSF for 7 days, culture media was changed every 3$^{rd}$ day with fresh MCSF. After 7-day culture, CD14 cells were harvested and cultured with hMSC in 1:1 ratio for 48 hrs. Reaction supernatant was collected to estimate $PGE_2$ concentrations and cells were processed to isolate total RNA for gene expression analysis.

### hMSC treatment in SAMP

SAMP-1/YitFc mice with established inflammation (>25 weeks) were used for the in vitro and in vivo studies. In vitro cultured hMSC ($5.0x10^6$) were resuspended in 200µl PBS and intraperitoneally injected in mice. An equivalent dose of 200 µl PBS served as vehicle control, while a daily dose of dexamethasone (2mg/kg of mice weight) for seven days was used as a positive control.

### Stereomicroscopic examination

The distal ileum (10 cm) was fixed in Bouin's solution and imaged using stereomicroscope (Amscope) with 10X optical zoom to score the inflamed areas. The tissue was imaged in 1 cm sections followed by a complete image compilation. The area with cobble stoning and villus flattening was considered inflamed, and the per cent disease area was calculated using the ImageJ software package.

### Tissue collection and Histological Evaluation

The distal ileum (10 cm) was fixed in Bouin's solution, paraffin-embedded, cut into 5µm sections and stained with hematoxylin/eosin. Histological assessment of tissues was performed by trained pathologist, blinded to the treatment. A standardized semi-quantitative scoring system was followed as described in the study.[4] The scoring system involve evaluation of 5 histologic parameters: (1) Villus

Architecture (2) active inflammation; and (3) chronic inflammation (4) mononuclear infiltration (5) Transmural inflammation.

**Gene Expression Analysis**

Total RNA was extracted using RNeasy Mini Kit, following the manufacturer protocol, and quantified using broad range Qubit 2.0 RNA Quantitation fluorometer kit. cDNA was synthesized using High-Capacity cDNA Reverse Transcription Kit with RNase Inhibitor. The primers were selected from the reported scientific literature; Oligonucleotide sequences are shown in Table below. Real-time qPCR was performed in Bio-Rad thermocycler (Bio-Rad C1000 CFX384) using SsoAdvanced Universal SYBR Green Supermix from BioRad. GAPDH used as the reference gene in the mRNA expression. The relative expression was obtained with BioradCFX Maestro 1.0 software and gene expression was determined using the standard curve method for each gene. The data are plotted as fold change using GraphPad Prism Version 9.0 (GraphPad Software, San Diego, CA). Genes primers sequence are provided in Table S1.

**Flow cytometry**

The mice were euthanized after the specified treatment period and MLN were harvested in the complete T cell media (RPMI1640+10% FBS+2mM Glutamine + HEPES + NAA + Sodium pyruvate + β-mercaptoethanol). The MLN were processed to form single cell suspension using the syringe top and 70-micron strainer. Cells were harvested and counted using a hemocytometer. $1x10^6$ cells were first incubated with the Fc block (anti-CD16/32), and later surface staining was performed using fluorochrome-conjugated monoclonal antibodies purchased from BioLegend: APC-anti-CD45 (30-F11), BV785-anti-CD3(17A2), BV605-anti-CD8α (53-6.7). Live/Dead$^{TM}$ fixable aqua dead cell stain kit

(Invitrogen) was used to stain the dead cells. Flow cytometry data were acquired on LSR Fortessa flow cytometer (BD Biosciences) and analyzed using FlowJo 10.6.1 software (FlowJo, LLC).

**In vivo phagocytosis by CD11b macrophages:**

In-vitro cultured hMSCs were trypsinized and stained with CellTracker$^{TM}$ Red CMTPX dye as per the manufacturer's protocol and injected peritoneal (5 million per mouse). Peritoneal lavage (P.L.) was performed on day 9 after injecting 10ml of PBS in peritoneal cavity, peritoneal fluid was centrifuged at 300 g for 5min to collect the cells. SVF tissue was processed as per the previously reported method by Hearnden et al.[5] Single cell suspension from P.L. and SVF was later stained for APC/Cy7 anti CD11b monoclonal antibody (BioLegend, M1/70).

**In vitro phagocytosis by peritoneal macrophages:**

Peritoneal lavage was performed in SAMP mice and peritoneal cells were collected after centrifugation at 300g for 5 min. Collected peritoneal cells were further enriched for peritoneal macrophages using MACS isolation kit. Enriched population of macrophages were cultured on glass cover slip in 6 well plates overnight in RPMI1640 medium. Live and apoptotic hMSCs were co-cultured with surface adhered macrophages in 1:1 ratio; hMSC were heat treated at 60°C for 45 minutes to induce apoptosis and imaged at different time interval to observe phagocytosis using brightfield microscope. For the fluorescence imaging, apoptotic hMSC were stained with annexin Pacific Blue (BioLegend) dye as per the manufacturer protocol and co-cultured with peritoneal macrophages for 2 hrs. After 2 hrs, cultured wells were washed thrice with PBS, fixed using 4% formaldehyde for 20 min at 4°C and permeabilized using 0.1% triton X100 for 15 min at room temperature (RT). Cells were further blocked for 30 min at RT using 1% BSA in PBS containing 0.1% tritonX100 and stained with PE/Cy-7 anti-F4/80 (BioLegend, BM8) as per manufacturer protocol. Cells were washed with PBS; glass cover slip were mounted on glass slides using 80% Glycerol in PBS as mounting media and imaged using fluorescence microscope (ZEISS).

**In vivo animal imaging:**

Bioluminescent imaging was performed after the subcutaneous (SQ) injection of 150 mg/kg of luciferin substrate, using a Xenogen IVIS 200 series system. Fifteen minutes after intraperitoneal administration of bioluminescent hMSCs, an early BLI is performed to evaluate cell distribution throughout the mouse body. Later, images are taken every alternate day to evaluate cell localization. Additionally, hMSC were tagged using a NIR dye IVISense680 following manufacturer protocol, and time course epifluorescence imaging was performed using IVIS Spectrum (Perkin Elmer). Following removal of abdominal hair and verification of lack of fluorescence from intestinal contents, $5x10^6$ labelled hMSC were peritoneally injected and live imaging was performed immediately day 0 and followed on days nine and 28. After completion of the experiment, colon, mLN tissue were harvested and single cell suspension was prepared to detect hMSCs using flow cytometery. Cells were stained using PE anti-human CD73(AD2), APC anti-human CD105 (SN6) antibodies to detect hMSC.

**MR imaging:**

Twenty-six SAMP mice (34-40 wk) with established SI inflammation together with parental AKR controls mice were imaged at the time of euthanasia in a 9.4 Tesla preclinical MRI scanner using the T2 fat suppression protocol in two independent experiments. To ameliorate SI inflammation, mice were injected with Dexamethasone 2mg/gram/daily intra-peritoneally (i.p) for 7 days prior to euthanasia, and eleven mice with 5 million human mesenchymal stem cells (hMSC) i.p. 4 weeks prior to euthanasia. As disease control, mice were treated with PBS i.p. four weeks prior to euthanasia. A radiologist blinded to treatment annotated the small bowel in mice. Histopathology was performed on small intestine samples of all the mice to correlate disease severity with MRI scores. Radiomic analysis was conducted within annotated bowel wall regions on the four axial sections. From each section, 100 radiomic features from 4 different classes were extracted to quantify bowel wall appearance via different texture and image analytic operators to yield quantitative measurements of image heterogeneity and gradient responses. Statistical feature selection and pruning was conducted to identify the top radiomic features that could best differentiate between groups selected via statistical testing. Top-ranked radiomic features primarily comprised local neighbourhood heterogeneity ($P<0.001$) and filtered intensity responses ($P<0.001$). These top-ranked features were used to train a machine learning classifier to yield a radiomics-based likelihood of disease severity for all SAMP mice (n=26), that was scaled to range from 0 to 20. A Small Intestinal MRI Pathology Inflammation Score was computed for each mouse based on the summation of radiomic and histology scores.

**Single cell RNA sequencing**

Stromal vascular fraction (SVF) of mice mesentery was isolated using a previously reported method by Hearnden et al.[5] Briefly, mice were euthanized after completion of treatment and mesenteries were precisely harvested in cold PBS. Mesenteric tissue was minced into small pieces and vortexed to collect SVF. SVF further subjected to enzymatic digestion at 37°C and 200 rpm for 30 min. using digestion media (Collagenase-Type A, Deoxyribonuclease 1, 100mM Calcium chloride; sucrose, BSA, HEPES, PBS). Enzymatic digest was centrifuged at 300g for 30 min at 4°C and pellet was resuspended in complete RPMI media and filtered through 70micron strainer. Single cell RNA sequencing was performed using SPLiTseq method from Parse Biosciences; cells were sequenced using NovaSeq PE100 illumine platform at the sequencing depth of 50000 reads/cells. Log fold change >1.5 and FDR<0.05 considered statistically significant in differentially expressed genes (DEGs).

**cDNA library preparation, sequencing, and data analysis**

Isolated single cells were counted using hemocytometer and fixed using cell fixation kit from Parse Biosciences. Fixed cells were counted using hemocytometer before cDNA library preparation and library construction was completed using Single Cell Whole Transcriptome-100k cells/nuclei kit from Parse Biosciences. 80,000 cells across 8 mice samples were initially fixed using cell fixation kit and later subjected to c-DNA library preparation. In total ~50,000 cells were recovered after fixation and 5 sub-libraries were prepared. Two sub libraries with ~20000 cells were sequenced at 50k read. Cells with fewer than 200 genes, with mitochondrial gene expression greater than 20%, and with number of UMIs below 200 or above 50000 were filtered. After completion of sequencing, raw sequencing data was deconvoluted to generate gene expression data for single cells, using Parse Biosciences' data analysis pipeline (split-pipe v0.9.3) with default parameters and GRCm39 mouse reference genome. Gene expression data were preprocessed, clustered, and UMAP coordinates were calculated using Seurat, version 4.1.0,[6] running in R version 4.1.0.[7] Data were log normalized and corrected prior to clustering by regressing out number of UMIs, percentage of mitochondrial gene expression, and cell cycle scores, using Seurat's ScaleData function. Clustering resolutions of 0.4 and 1 were chosen to present cluster in day 9 and day 28 samples. Differential expression (DE) analyses were conducted on log normalized data using limma, version 3.50.3.[8] The model used in limma included effects for treatment, sample within treatment, and percent mitochondrial gene expression. DE analyses included genes expressed in at least 10% of cells out of all cells of a given type or included in each analysis.

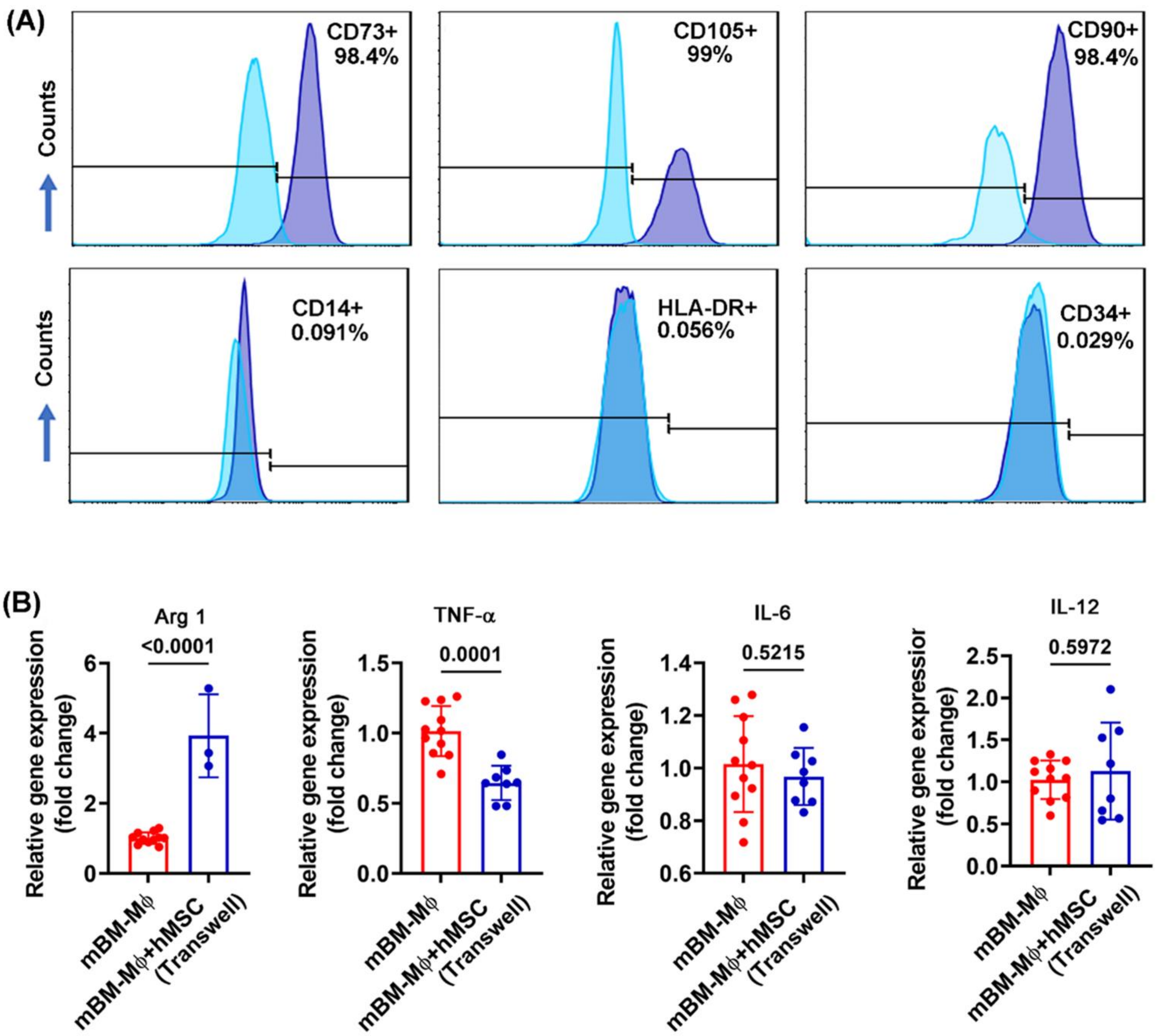


**Figure S1.** Human bone marrow mesenchymal stem cells characterized by **(A)** positive CD73, CD105, CD90 and negative CD14, HLA-DR, and CD34 surface marker **(B)** Relative gene expression of indicated markers and cytokine measured in total RNA extracted from murine bone marrow-derived macrophages co-cultured with hMSC (separated through trans well). The gene expression was determined by qRT-PCR, normalized to GAPDH, and expressed as fold change (2-ΔΔCt); Data correspond to two independent experiments, and each data point represents one mouse.

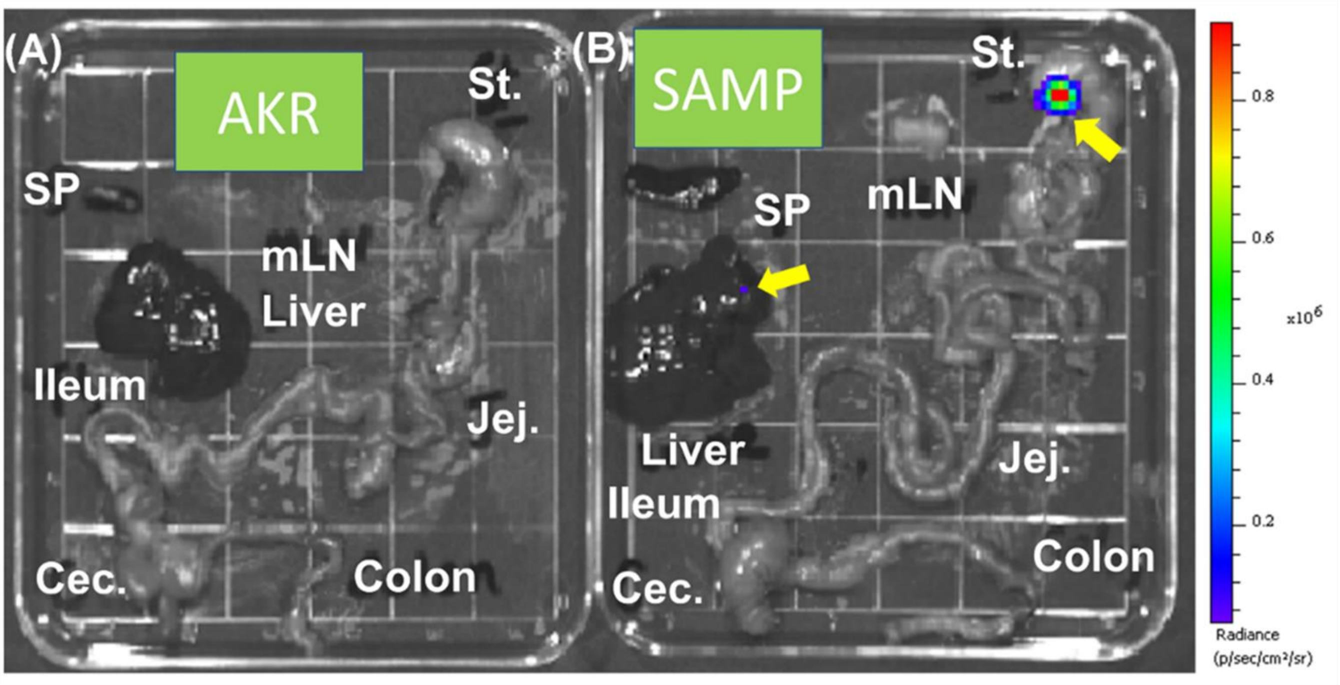


**Figure S2:** Ex vivo bioluminescent imaging performed on harvested body organs showing the representative image of **(A)** AKR mice and **(B)** SAMP mice. SAMP mice show the presence of live hMSCs in the liver and the proximity of the stomach. (SP=spleen, St=Stomach, mLN=mesenteric lymph node, Jej.=Jejunum, Ileum, Cec.=Cecum). Bold arrow showing the presence of bioluminescent hMSC.

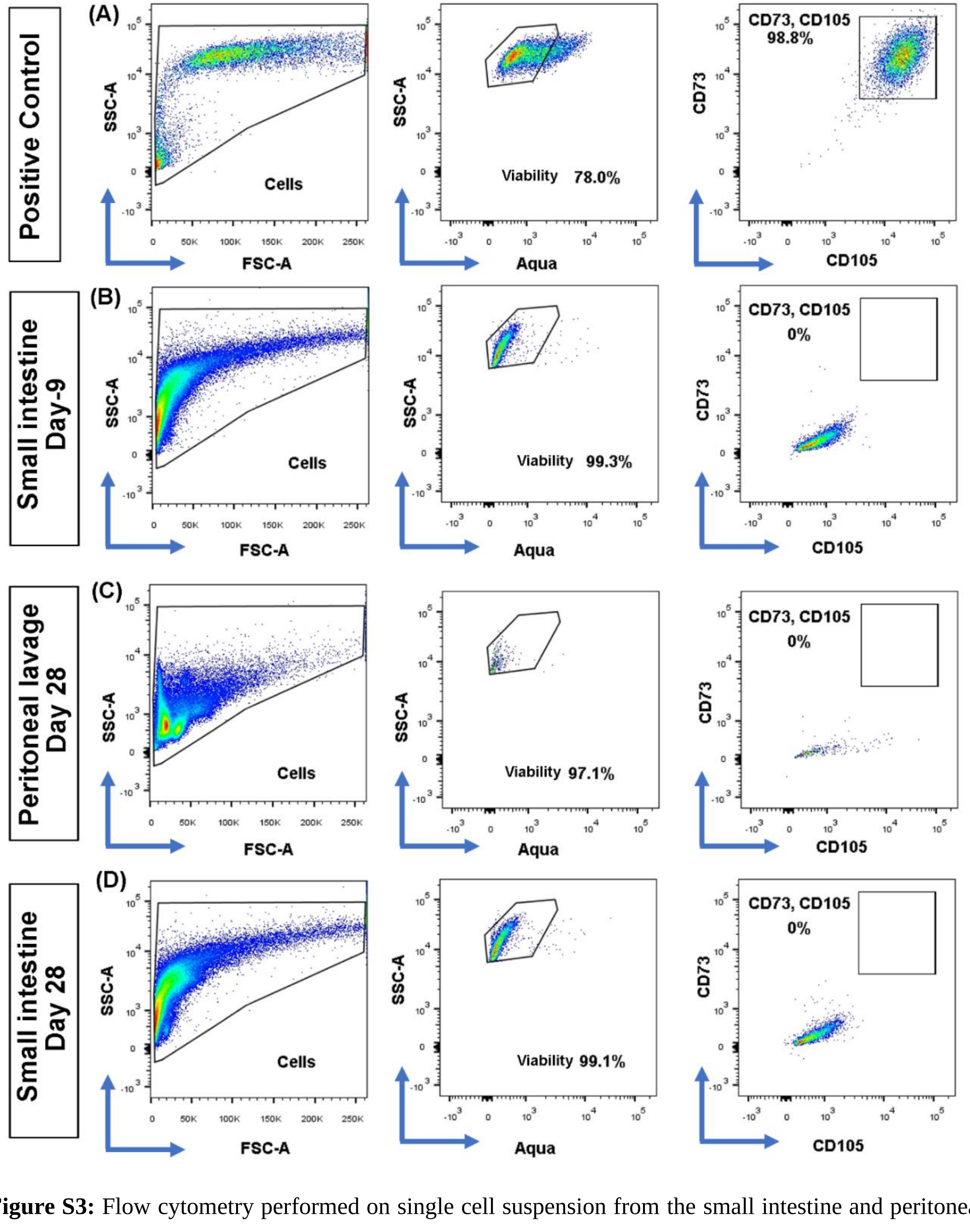


**Figure S3:** Flow cytometry performed on single cell suspension from the small intestine and peritoneal lavage cell isolated at day 9 and day 28 after peritoneal administration of 5 million IVISense680 tagged hMSC. Gating scheme showing hMSC ($CD73^+$, $CD105^+$) population. **(A)** In vitro cultured hMSC as a positive control, **(B)** Single-cell suspension of the small intestine, on day 9, **(C)** Peritoneal lavage on day 28, **(D)** Single-cell suspension of the small intestine, on day 9. Data represent 3 biological replicates.

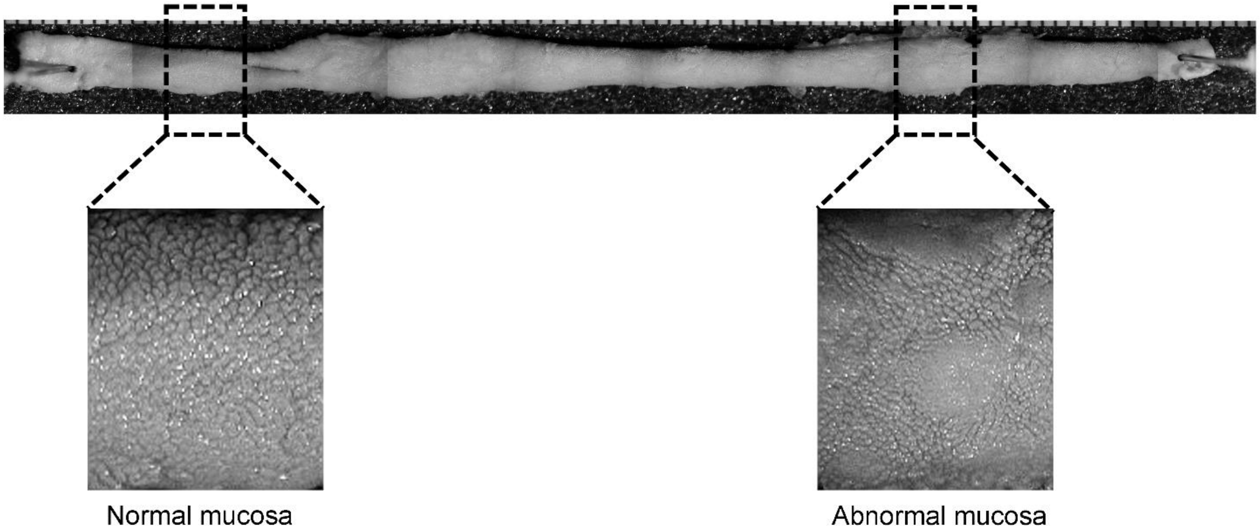


**Figure S4.** Representative stereomicroscopic image of SAMP small instestine showing difference between normal and abnormal mucosa.

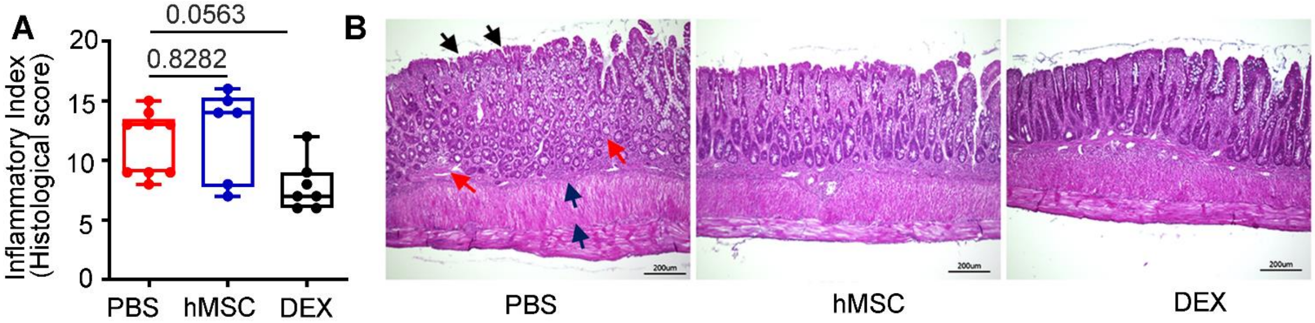


**Figure S5** SAMP mice treated with hMSC, and DEX did not achieve histological healing by day 9 **(A)** Inflammatory Index for disease severity is a histological index composed of the villus distortion index, active inflammation index, mononuclear inflammation index, chronic inflammation index and transmural inflammation index. SAMP mice treated with hMSC, and DEX did not achieve significant histological healing (P=0.8282), however, a partial response is observed in DEX-treated SAMP (P=0.0563). (**B**) Representative histopathology photomicrograph of ileum tissue from PBS, hMSC and DEX-treated groups. PBS-treated group show histological features of villus distortion (black arrow), Immune infiltrates and crypt hyperplasia (red arrow), and muscle hypertrophy (blue arrow). The scale bar represents 200µm.

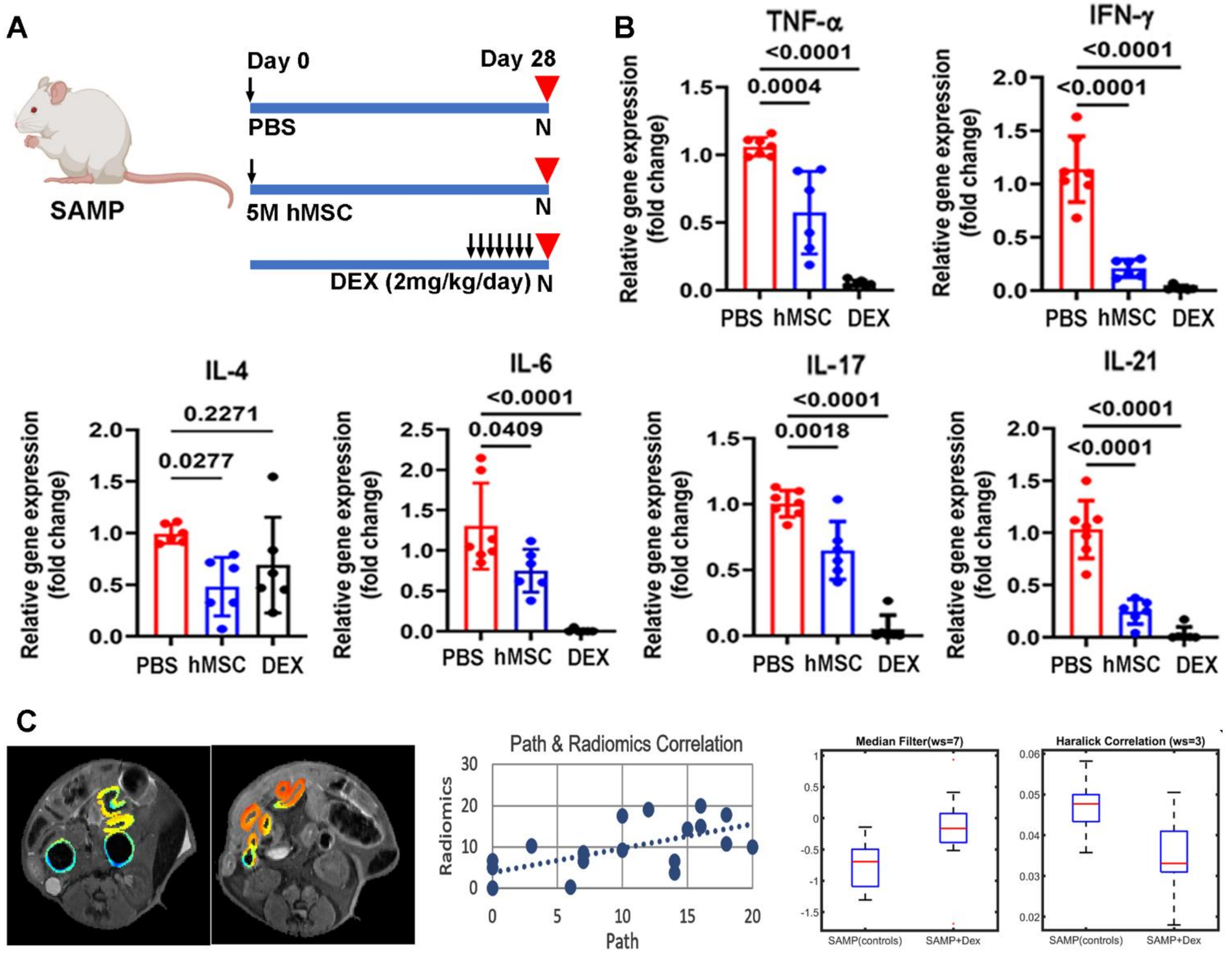


**Figure S6: (A)** Schematic showing treatment regime of the experiment **(B)** Relative gene expression of Th-1, Th-2, and Th-17 pathway cytokines measured in total RNA extracted from mLN cells. The gene expression was determined by qRT-PCR, normalized to GAPDH, and expressed as fold change (2-ΔΔCt). hMSC-treated mice showed a decrease in gene expression of pro-inflammatory Th-1 cytokines TNF-α (20.57-fold change, P<0.001) and IFN-γ (0.21-fold change, P<0.0001); IL-6 (0.75-fold change, P<0.05) and Th-17 cytokine IL-17(0.65-fold change, P<0.01). DEX significantly downregulated gene expression of the majority of proinflammatory cytokines including TNF-α (0.05-fold change, P<0.0001) IFN-γ (0.02-fold change, P<0.0001), IL-6 (0.01-fold change, P<0.0001), IL-17(0.05-fold change, P<0.0001). 6-8 mice/group were studied in 3 independent experiments. **(C)** Workflow showing the statistical evaluation from MR Images; feature extraction, correlation between path and radiomics scores, top radiomic features from SAMP mice selected via statistical testing.

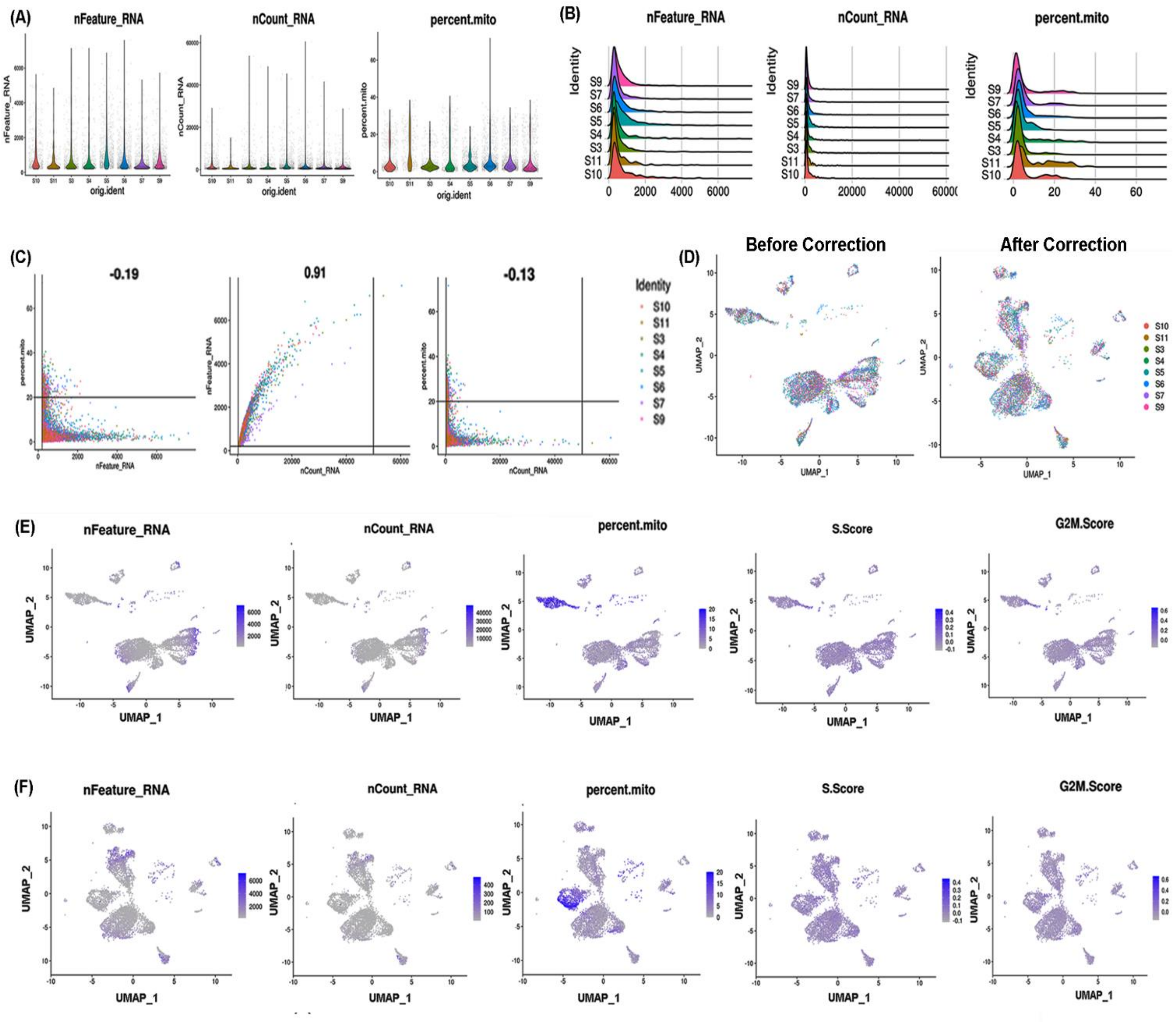

**Figure S7.** Single-cell RNA sequencing data pre-processed using Seurat, version 4.1.0 running in R version 4.1.0 for quality control metrics showing **(A)**Violin plots **(B)** Ridge plots **(C)** scatter plots with proposed cutoffs **(D)** UMAP plot by sample, prior to correction and after correction **(E)** UMAP plot showing the number of UMIs/cell, number of genes/cell, percent mitochondrial gene expression, cell cycle S score, cell cycle G2M score before correction and **(F)** after correction.

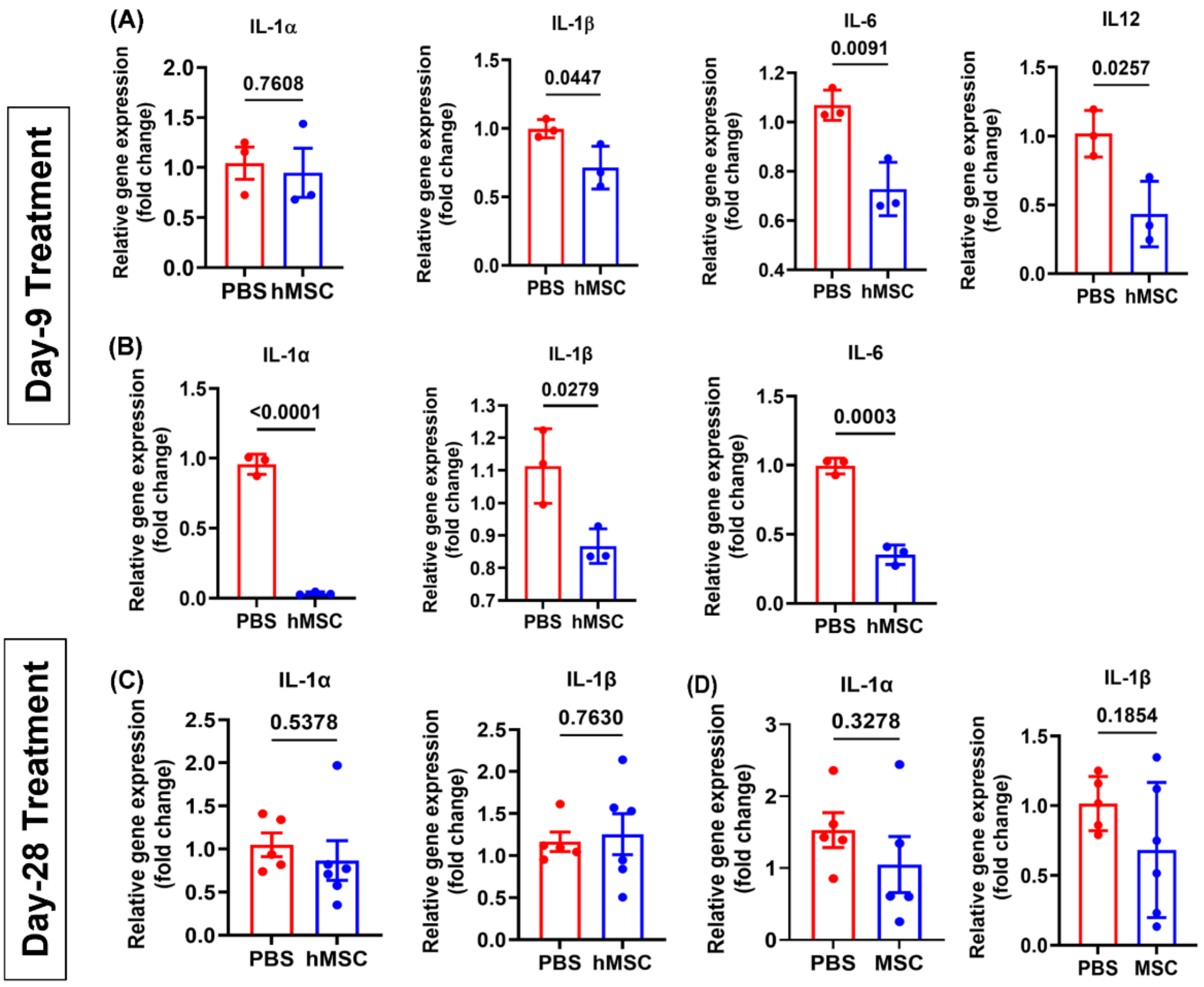


**Figure S8.** Relative gene expression of cytokines measured in total RNA **(A)** hMSC treatment at day 9, resulted in the anti-inflammatory phenotype of mLN CD11b$^+$ macrophages with downregulation of IL-1β (0.71-fold change, P=0.044), IL-6 (0.71-fold change, P=0.0091) and IL-12 (0.43-fold change, P=0.025), while no change in gene expression was observed for IL-1α (P=0.76). **(B)** CD11b$^+$ macrophages isolated from SVF of mesentery also demonstrated similar anti-inflammatory phenotype with downregulation of pro-inflammatory cytokines IL-1α (0.03-fold change, P<0.0001), IL-1β (0.86-

fold change, P=0.027), and IL-6 (P=0.0003). Relative gene expression of IL-1α and IL-1β did not change significantly at day 28 in **(C)** mLN and **(D)** SVF-derived CD11b$^+$ macrophages. The gene expression was determined by qRT-PCR, normalized to GAPDH, and expressed as fold change (2-ΔΔCt). Data correspond to two independent experiments and each data point represents one mouse.

### Table S1: RT-qPCR primers sequence

| **Gene** | **Forward (‘5-3’)** | **Reverse (‘5-3’)** |
|---|---|---|
| GAPDH | CCCATCACCATCTTCCAGGAGC | CCAGTGAGCTTCCCGTTCAGC |
| TNF-α | GCCTCTTCTCATTCCTGCTTG | CTGATGAGAGGGAGGCCATT |
| IL-12p40 | CAGAAGCTAACCATCTCCTGGTTTG | TCCGGAGTAATTTGGTG CTTCACAC |
| IL1-α | TCTCAGATTCACAACTGTTCGTG | AGAAAATGAGGTCGGTCTCACTA |
| IL1- β | CCTTCCAGGATGAGGACATGA | TGAGTCACAGAGGATGGGCTC |
| Arginase I | CAGAAGAATGGAAGAGTCAG | CAGATATGCAGGGAGTCACC |
| IL-6 | GAGGATACCACTCCCAACAGACC | AGTGCATCATCGTTGTTCATACA |
| IFN-γ | CGACTCCTTTTCCGCTTCCTGAG | TGAACGCTACACACTGCATCTTGG |
| IL-4 | AGATGGATGTGCCAAACGTCCTCA | AATATGCGAAGCACCTTGGAAGCC |
| IL-17 | ATCCCTCAAAGCTCAGCGTGTC | GGGTCTTCATTGCGGTGGAGAG |
| IL-21 | TCAGCTCCACAAGATGTAAAGGG | GGGCCACGAGGTCAATGAT |

### REFERENCES


1. Lennon DP, Caplan AI. Isolation of human marrow-derived mesenchymal stem cells. *Experimental hematology* **34**, 1604-1605 (2006).

2. Fierro FA, *et al.* Mesenchymal stem/stromal cells genetically engineered to produce vascular endothelial growth factor for revascularization in wound healing and ischemic conditions. *Transfusion* **59**, 893-897 (2019).

3. Sorrell JM, Baber MA, Caplan AI. Clonal characterization of fibroblasts in the superficial layer of the adult human dermis. *Cell and tissue research* **327**, 499-510 (2007).

4. Burns RC, Rivera-Nieves J, Moskaluk CA, Matsumoto S, Cominelli F, Ley K. Antibody blockade of ICAM-1 and VCAM-1 ameliorates inflammation in the SAMP-1/Yit adoptive transfer model of Crohn's disease in mice. *Gastroenterology* **121**, 1428-1436 (2001).

5. Hearnden R, Sandhar B, Vyas V, Longhi MP. Isolation of stromal vascular fraction cell suspensions from mouse and human adipose tissues for downstream applications. *STAR Protoc* **2**, 100422 (2021).

6. R Core Team (2021) R: A language and environment for statistical computing. *R Foundation for Statistical Computing, Vienna, Austria* (2021).

7. Hao Y, *et al.* Integrated analysis of multimodal single-cell data. *Cell* **184**, 3573-3587 e3529 (2021).

8. Ritchie ME, *et al.* limma powers differential expression analyses for RNA-sequencing and microarray studies. *Nucleic acids research* **43**, e47 (2015).
